# An Optimization Framework for the Design of Radiofrequency Coils for Magnetic Resonance Imaging

**DOI:** 10.1101/2025.08.04.668545

**Authors:** José E. Cruz Serrallés, Ilias I. Giannakopoulos, Siqi Wang, Damien Chen, Daniel Zint, Daniele Panozzo, Denis Zorin, Riccardo Lattanzi

## Abstract

The radiative characteristics of the radiofrequency receive coils dictate the signal-to-noise ratio (SNR) of magnetic resonance images. Despite the crucial importance of RF coils, the practical coil design process has remained a largely empirical one. This work introduces a novel optimization framework for rational coil design, which relies on a fully automated pipeline that combines rapid electromagnetic simulations, shape optimization and coil meshing. The objective function iteratively maximizes SNR performance in a target region of interest with respect to the ultimate intrinsic SNR, which is the theoretically highest SNR independent from any particular coil design. The forward simulation employs a fast electromagnetic solver based on coupled surface and volume integral equations. The coils are represented as B-spline curves with an associated width, and automatically meshed for EM simulation. We implemented a new method to tune and decouple coils at each iteration without manual user intervention. The algorithm optimizes the size and position of a given number of coils with a combination of grid search and a line search. We demonstrated the framework by designing receive arrays of increasing complexity that yield optimal SNR for different target regions inside a numerical head model. SNR simulation time ranged from 15 s for a 3-coil configuration to 32 s for a 12-coil array, constrained to a helmet-like surface, including tuning and decoupling. The optimized 12-coil geometry yielded 9% higher average SNR performance in the brain at 3 T. This work represents the first automated coil optimization framework that uses full-wave electromagnetic simulations and ultimate performance benchmarks. This novel approach enables the systematic design of coils for magnetic resonance imaging with significantly improved SNR performance, potentially transforming coil development from empirical design to physics-driven optimization.

## 1 Introduction

Radiofrequency (RF) coils are at the heart of any magnetic resonance (MR) imaging application. They are the source of the RF fields that generate MR signals and the mechanism by which RF fields generated by excited nuclear spins are detected. RF coil design is therefore a critical determinant of the performance of all MR imaging (MRI) systems. As the number of channels available in MR systems has increased to enable faster acquisitions with parallel MRI [1–3], building prototypes of coil arrays has become more difficult and expensive; as a consequence, coil design has relied ever more on electromagnetic (EM) simulations. While careful coil design is important at all field strengths, appropriate coil designs are truly essential for high-field MRI, both for the preservation/improvement of image quality and for the avoidance of adverse effects in patients. This work aims to introduce a novel optimization framework for rational RF coil design.

For receive arrays, the number, geometry, and arrangement of each coil element, as well as interelement decoupling, coil loading, and parallel imaging performance, are important design parameters that can impact the overall signal-to-noise ratio (SNR) [1–3]. However, current approaches to coil design (Section 2.1 are limited by time-consuming simulations and a lack of systematic, automated tools to explore complex design spaces.

This previous work does not address another major limitation of the current approach to coil design: the quality of a coil is typically judged in comparison to other available coils, giving no indication of whether there is room for further improvement beyond the best-performing design tested. To address this, theoretical coil performance limits [4–7], such as the ultimate intrinsic SNR (UISNR), could be used as absolute references during coil design [7–11]. The UISNR is the highest possible SNR compatible with electrodynamics and independent from any particular coil design [5, 12–14]. The UISNR represents the highest SNR allowed by electrodynamics, independent of any specific coil configuration. Prior studies (Section ?? showed that even state-of-the-art coils capture only a fraction of this limit in cortical regions, especially at ultra-high field strengths, leaving substantial room for improvement. This suggests the need for alternative design strategies that aim to approach ultimate performance.

UISNR integration into the coil design process has remained an open problem. To address this, we employed the MRGF concept to develop a novel shape optimization technique that automatically enhances coil configurations based on ultimate performance benchmarks. Given an existing coil configuration, we compute its variations for a set of design parameters to find a design that improves coil performance. This approach has been highly successful in material design, structural mechanics, and fluid dynamics, but has yet to be effectively applied to MRI coil design [15–19].

A key challenge is the need for an unconditionally robust simulation and modeling pipeline. At each step of the optimization, a new coil geometry must be generated, meshed, tuned, and simulated with no human intervention – a task currently infeasible with standard MRI simulation tools, in which users routinely spend days setting up a single simulation [20].

We propose an integrated approach to tackle this problem, which jointly considers forward simulation, shape computation, and coil meshing. Together with a new framework for rational coil design based on shape optimization, this work introduces several innovations: an accurate and fast solver for the surface integral equation, a method to automatically tune RF coils, an approach to mimic preamplifier decoupling in coil simulations, and a method for automatic meshing of coil geometries.

We demonstrate the use of our optimization framework through numerical examples of increasing complexity, from single-coil positioning, to shape optimization of small arrays, to large-array performance optimization toward the UISNR. We also show that the optimization results are consistent for different anatomical models.

The rest of the manuscript is organized as follows. In Section 2, we present a brief overview of previous work that is relevant to this paper. In Section 3, we specify the constraints of the design space for our proposed coil optimization. In Section 4, we describe our approach to efficiently model RF coils, whereas in Section 5, we describe the new SIE solver and the proposed method for automatic coil tuning and ideal decoupling. In Section 6, we introduce the coil design optimization algorithm and in Section 7, we provide details of the numerical experiments performed in this study to demonstrate it. The results are presented in Section 8 and discussed in Section 9, whereas Section 10 summarizes the main points of this work.

## 2 Related Work

In this section we briefly review related work in MRI coil optimization and shape optimization. Some more in-depth discussion of most closely related numerical methods can be found in Section 5.1.4.

### 2.1 Coil design optimization

Initial work on coil design optimization was performed in the quasi-static regime due to lower field strengths and hence lower operating frequencies [21–25]. Analytical methods, such as spherical harmonics [21] and cylindrical harmonics (Fourier-Bessel series) [22], were employed for the design of receive coils, gradient coils, and solenoidal coils. Analytical methods were popular because their corresponding basis and testing functions yield diagonalized systems, which are easily invertible. Later on, the quasistatic Biot-Savart Law was used explicitly to design gradient and RF coils, along with an error function that was minimized using conjugate gradient descent [23]. Further work involved variations of these basic approaches, such as randomized Monte Carlo sampling to match desired spherical harmonics [24], applications of the Biot-Savart Law to obtain desired field patterns (inverse design), and full-wave simulation using thin-wire approximations [25], among others. Quasi-static coil design methods have remained popular tools, especially for gradient coils, since these operate at much lower frequencies than RF coils. More recent examples of quasi-static coil optimization include parametric optimization of 1- and 2-loop arrays using the Biot-Savart law [26], boundary element method-based field profile matching with ohmic loss penalties for gradient coil design [27], and multi-objective field matching for gradient and shim coil design [28], among others [29].

Biot-Savart and quasi-static harmonic solutions are no longer accurate at MRI field strengths higher than 1.5 T. While Jefimenko’s Equation [30] offers a full-wave equivalent of the Biot-Savart Law, modeling using this equation can be difficult due to the strong 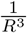 singularity in the Green’s function and requires making approximations that might not be valid, such as uniform current distribution in a loop. Early attempts at coil optimization in the full-wave regime include the application of the Helmholtz Green’s function with so-called stream functions to design a loop at 4.5 T [31], the design of a body coil by simulating first with Biot-Savart and then with Method of Moments discretization of the EFIE formulation while optimizing with a genetic algorithms metrics such as slice B_1_ homogeneity [32], and the application of a genetic algorithm to body coil design using a method-of-moments approach [33]. More recent examples in the full-wave regime include an inverse design approach for the design of a uniplanar RF coil while optimizing for RF field homogeneity [34] and parametric coil optimization for the design of RF transmit arrays while optimizing for slice homogeneity [35].

Another work solved an optimization problem to find the current density that maximized SNR for parallel imaging applications [36]. All these works focused on the design of one- or two-element arrays, rather than dense arrays, and used simple geometries to mimic the anatomy. Genetic algorithms have also been proposed to simultaneously optimize multiple design parameters for birdcage coils or two-element arrays, also in this case using uniform objects with simple geometries [17–19]. The main limitation of these previous studies was the time-consuming EM simulations that prevented the implementation of iterative optimization algorithms to design many-element arrays using realistic anatomical models.

The Magnetic Resonance Green Function (MRGF) method, based on a fast EM solver tailored to MRI applications, was more recently proposed to simulate in minutes the EM field inside a realistic anatomical model for any RF coil designed over a specified substrate for which the MRGF had been precomputed [37]. A proof-of-concept study suggested that the MRGF method could be employed to iteratively optimize size and position of two transmit coils to maximize magnetic field homogeneity inside a numerical human head model [35]. No other attempts have been made at using MRGF to automatically optimize the design of arrays with a larger number of coil elements. The UISNR mentioned in the intro provides an absolute benchmark for coil design, representing the highest possible SNR compatible with electrodynamics, and was considered in a few previous works [5–7, 12–14]. It is computed using a complete EM basis to simulate idealized, infinite arrays. Prior studies have shown that conventional coils achieve only a fraction of the UISNR, particularly in superficial cortical regions and at higher field strengths [4, 8–11]. This underscores the significant potential for improvement. More recently, UISNR has been used to directly inform coil design and optimization pipelines [20, 38–40], motivating the approach taken in this work.

### 2.2 Shape optimization

There is an extensive literature on shape optimization in a variety of contexts, ranging from fluid dynamics to electromagnetic modeling and metamaterial design. The general mathematical foundations of PDE-constrained shape/topology optimization can be found in classical references [41–44].

Most of these work focus on optimization of surfaces bounding 3D domains or planar curves bounding 2D domains in the context of fluid or heat flows or elasticity. A recent survey [45] focuses on shape optimization in the context of electromagnetic modeling. POE-constrained shape and topology is widely used for metamaterial design in the context of elasticity [46, 47], electromagnetics and optics [48, 49]. PDE-constrained optimization of curves *on surfaces* of the type we perform in our work is less common. Most of the work is related to either curve smoothing/fitting to data e.g., adaptation of active contours to surfaces [50] and other smoothing methods [51–53]. Our approach to avoiding overlaps in coils, based on the contact potential introduced in [54] and extended to codimensional objects in [55] is related to the potential-based repulsive curves approach considered in [56]. A constrained-optimization based approach to self-avoiding curves is applied to polymer adsorption on surfaces in [57].

## 3 Coil Design Constraints

The coil optimization pipeline must satisfy specific constraints to ensure that the array can be constructed as designed and utilized effectively for MRI. The first constraint is that a coil cannot intersect itself, and no two coils should touch each other. Otherwise, the current patterns would be modified, and for example, two touching coils would effectively become a single, larger coil. In addition, these topology changes could result in sharp changes in the objective function with a negative effect on the optimization algorithm. The second constraint is that the coils cannot be too small, otherwise the SNR would be dominated by electronic noise and the loading of the sample would be poor, resulting in low performance [58]. The third constraint is that the optimized values for lumped elements (capacitance, inductance, and resistance) should fall within a user-defined range that reflects commercially available capacitors. The fourth constraint is that the coils must belong to the parametrization domain of the surface of the substrate (Section 4.2). During the automated design process, the first constraint is guaranteed by selecting a design space that satisfies it by construction. While optimizing the coil geometry, this is achieved by enforcing a barrier potential term, where we consider (1) a coil edge and the parameterization domain boundary edge; (2) edges within one individual coil to be the set of possible contact pairs (see Figure 1 and Section 6.1). We also introduced bridges (Section 4.4) to avoid coils touching each other. Instead of specifying a minimum value for the coil radius, the second constraint is automatically enforced by adding coil noise into the SNR calculation. In this work, we used only capacitors as lumped elements, and we satisfied the third constraint by specifying the minimum and maximum capacitance to be 1 pF and 200 pF, respectively, in the input JSON. These values are read and enforced during tuning and ideal decoupling (Sections 5.4.1 and 5.4.2). The fourth constraint is enforced by construction during the meshing process (Section 4.4).

**Fig. 1:**
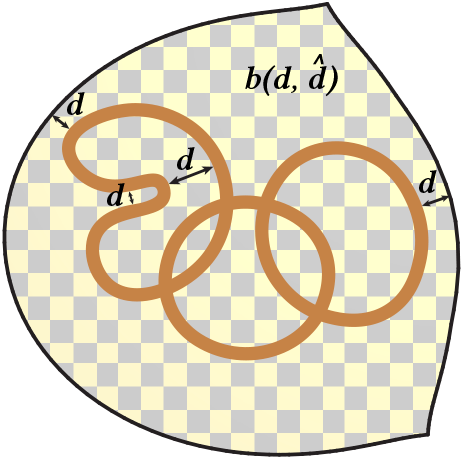
Coils are constrained so that they do not self intersect nor go outside the domain of the substrate. We implemented this by adding a barrier energy term 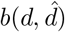, which vanishes if the distance *d* between contact pairs is larger than a predefined 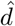 threshold.

## 4 Modeling

We represent coils in a parametrized way, suitable for optimization. The coil parametrization is established in two steps. First, we construct a map from a 2D domain to the 3D surface of the coil former (Section 4.2). Second, we represent coils as parametric curves with width in the 2D domain (Section 4.3). Finally, we discretize the parametrized coil design (Section 4.4). To facilitate interactive coil design, a graphical user interface (GUI) was developed (Section 4.1).

### 4.1 User interface

The GUI was implemented in TypeScript and based on the Babylon.js framework (Figure 2). The GUI allows users to place and adjust control points within a 2D domain, where coil geometries are rendered in real-time as a collection of B-splines. The interface supports intuitive translation, scaling, and duplication of entire coils, individual strips, or single control points. Lumped element components (Section 4.3) can be both visualized and parametrically adjusted through dedicated property panels. All modifications to the coil geometries are automatically reflected within the integrated 3D scene, which also allows users to overlay visualizations of volumetric body models that can be loaded as isosurfaces. A 3D cursor tool enables precise selection and transformation of coil elements within the 3D view, ensuring that users can configure coil geometries and their associated lumped element properties in a single, closed environment.

**Fig. 2:**
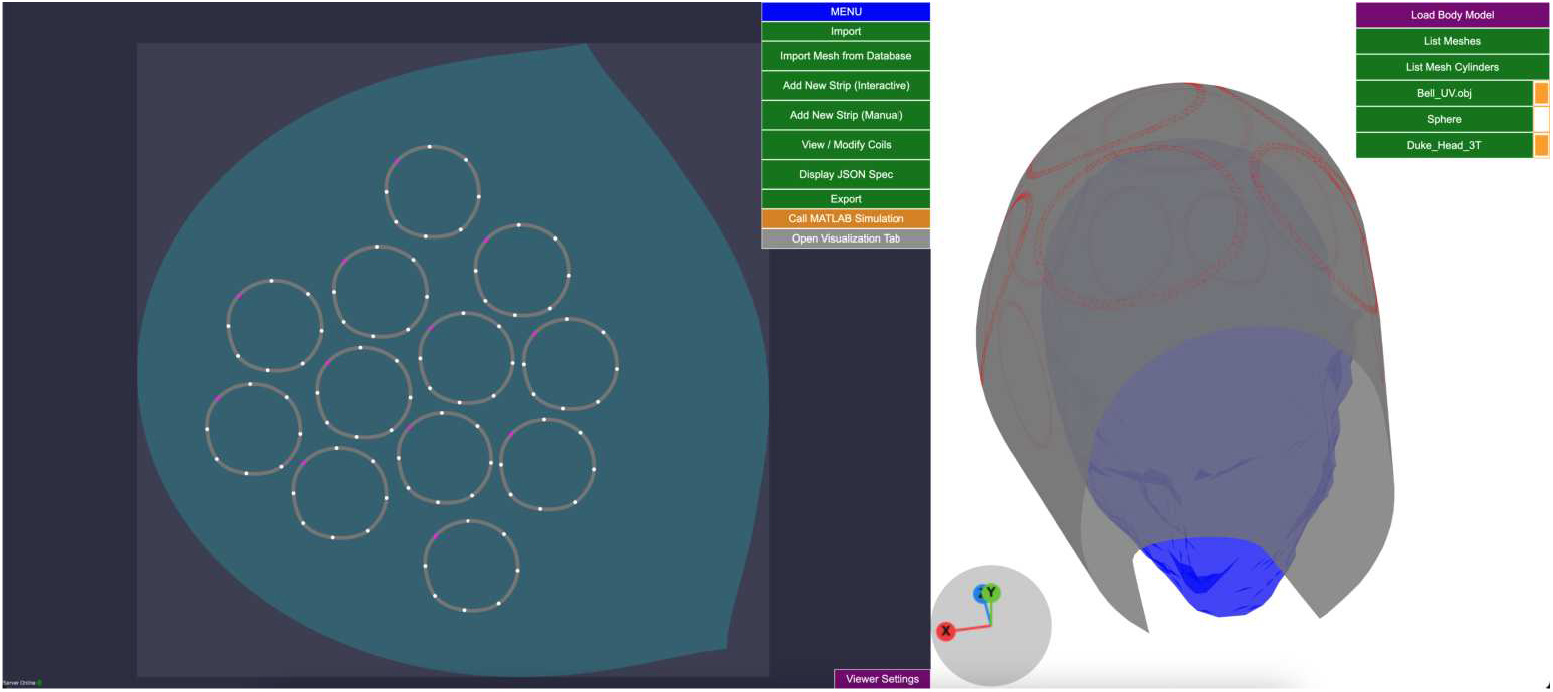
The graphical user interface (GUI) to manually design or display coils. The GUI visualizes coils and the substrate using both 2D and 3D interconnected views.

The parametrized coil design (control points, width, lumped elements information etc.) can be exported in a human-readable JSON file. This file is also the input for the optimization pipeline (Section 6.2).

### 4.2 Substrate parametrization

We assume that the substrate surface, i.e., the coil former, is given as a triangular surface mesh, which is a collection of triangles embedded in 3 dimensions. The triangular surface mesh is flattened in 2D with the method of Scalable Locally Injective Maps (SLIM) [59]. SLIM minimizes distortion energies with a re-weighting scheme and prevents flipped geometry. Any position u in 2D can be mapped to a position on the 3D substrate surface with Φ(u) : ℝ^2^ → ℝ^3^, by finding the nearest 2D triangle, computing the position’s barycentric coordinates for that triangle, and evaluating the coordinates in the corresponding 3D triangle (Figure 3). For efficient evaluation of Φ(u), we use an axis-aligned bounding box (AABB) tree data structure [60] to find the nearest triangle.

**Fig. 3:**
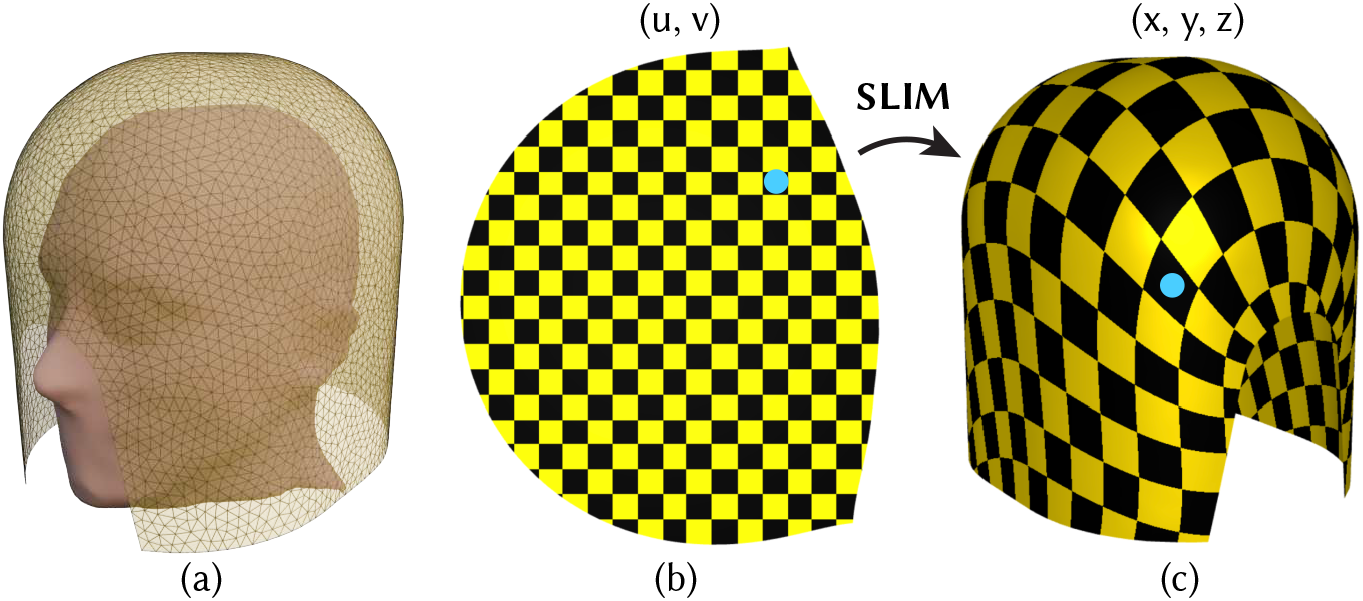
(a) Geometry of an example substrate surface mesh outside the head model. (b,c) Scalable Locally Injective Maps (SLIM) [Rabinovich 2016] with symmetric Dirichlet to minimize arbitrary distortion energies.

### 4.3 Coil parametrization

We use B-splines *B*(*t*), *t* ∈ (0, 1) as parametric curves to represent coils in 2D. Every coil has a width *w*, such that the entire coil domain in 2D is described as *B*(*t, s*), *t* ∈ (0, 1), *s* ∈ (−*w/*2, *w/*2), where ∇*s* is orthogonal to ∇*t*. The 3D coil geometry is described by Φ(*B*(*t, s*)).

RF coils have electrical components (capacitors, inductors, resistors, or feeding ports) attached to them that also need to be part of the coil parametrization. We represent those components as lumped elements and store their parametric position *t* ∈ [0, 1) within the B-spline to which they are attached. Note that lumped elements always span the entire coil width, i.e., *s* ∈ (−*w/*2, *w/*2).

### 4.4 Meshing

We discretize the coils in 2D with triangle meshes and then map those triangles to the 3D surface (Figure 4). The entire meshing pipeline consists of: (1) converting the B-splines into polylines, (2) generating strips from the polylines by offsetting them by *w/*2 in both directions, (3) inserting a line for each lumped element, (4) triangulating, and (5) mapping to 3D.

**Fig. 4:**
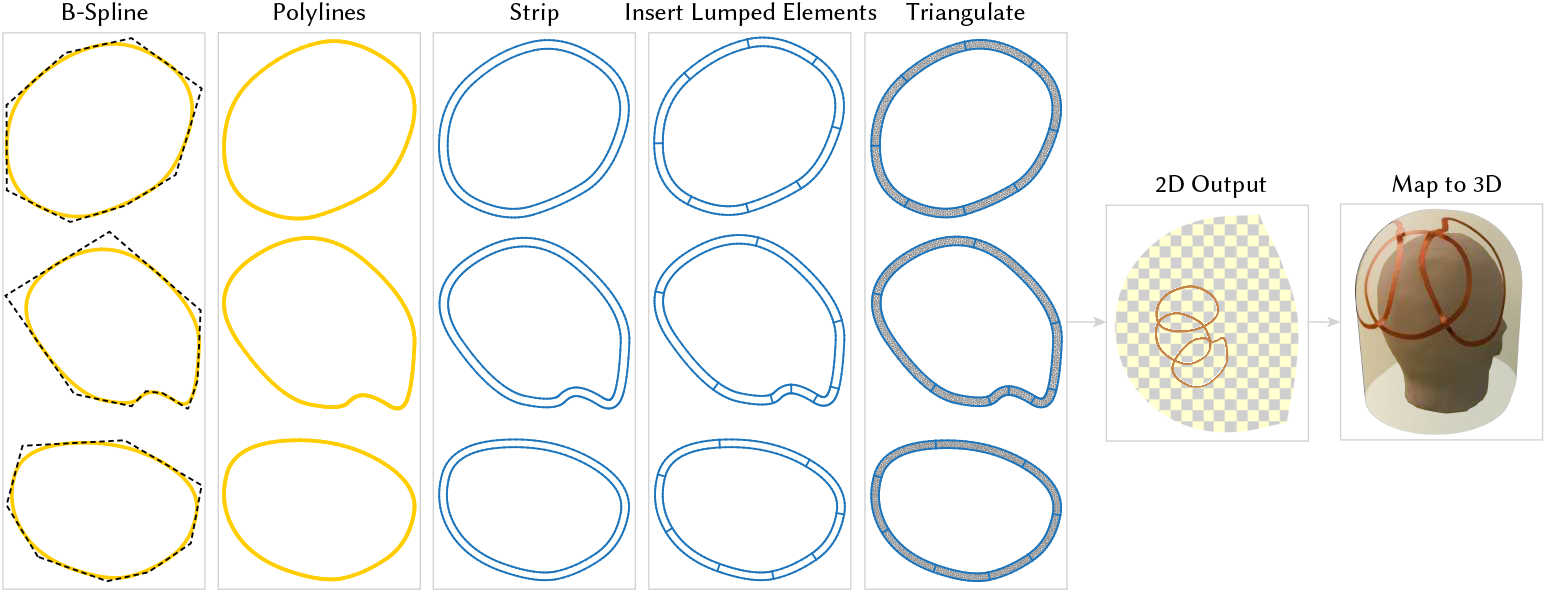
Overview of the meshing pipeline.

#### Conversion into polylines

For simpler handling, we convert the B-splines into Bézier curves. We recursively subdivide the Bézier curves at *t* = 0.5 until the flatness of each curve is below a certain threshold *τ* [61]. For a cubic Bézier curve with four control points [**b**_0_, **b**_1_, **b**_2_, **b**_3_], the flatness of a curve *b*(*t*) is defined as

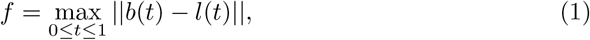

where *l*(*t*) is the line segment between the start point **b_0_** and end point **b_3_** (see Figure 5), which can also be represented as a cubic Bézier curve with control points [**b**_0_, (2**b**_0_ + **b**_3_)*/*3, (**b**_0_ + **2b**_3_)*/***3, b_3_**]. Hence we have

**Fig. 5:**
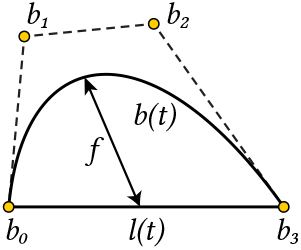
Illustration for the flatness evaluation in Equation (1), where the flatness *f* of a cubic Bézier curve *b*(*t*) is defined as the maximum distance between *b*(*t*) and *l*(*t*).

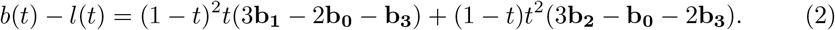

For *ξ* = (**3b_1_** − **2b_0_** − **b_3_**) and *η* = (**3b_2_** − **b_0_** − **2b_3_**), we obtain an upper bound for the flatness,

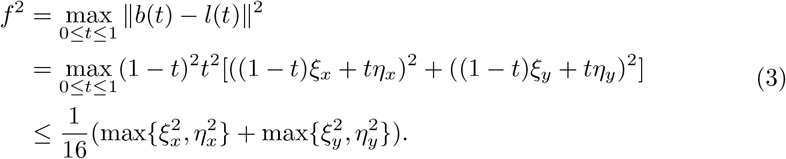

By recursively subdividing the curves until 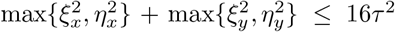 is satisfied, and replacing all curves with line segments, we guarantee to introduce a discretization error of at most *τ*.

#### Strip generation

We generate a strip contour from the polylines by offsetting each line segment by *w/*2 in the positive and negative normal direction and computing their intersections (Figure 6). The vertices in the interior of a strip are the intersections between two adjacent offset polylines; for example, in Figure 6, *A* and *B* are computed by intersecting lines *l*_1_ and *l*_2_, and *l*_2_ and *l*_3_, respectively. The polyline’s boundary vertices are offset in the normal direction of the incident line segment. In this way, we obtain the Planar Straight Line Graph (PSLG) of the strips. To avoid introducing small discretized elements due to short lines in the PSLG, we collapse edges shorter than a tolerance of *ϵ* = 10^*−*3^, starting with the shortest edge (Algorithm 1).

**Fig. 6:**
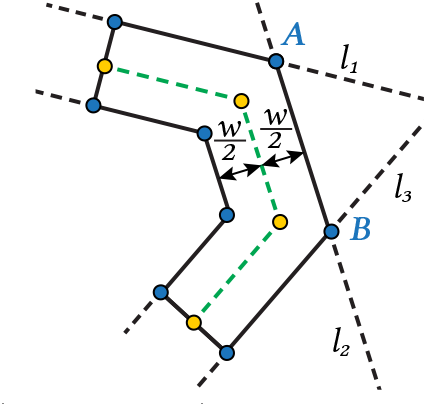
Strip generation (solid black) by offsetting polylines (dashed green).

##### Algorithm 1 PSLG decimation for strip generation

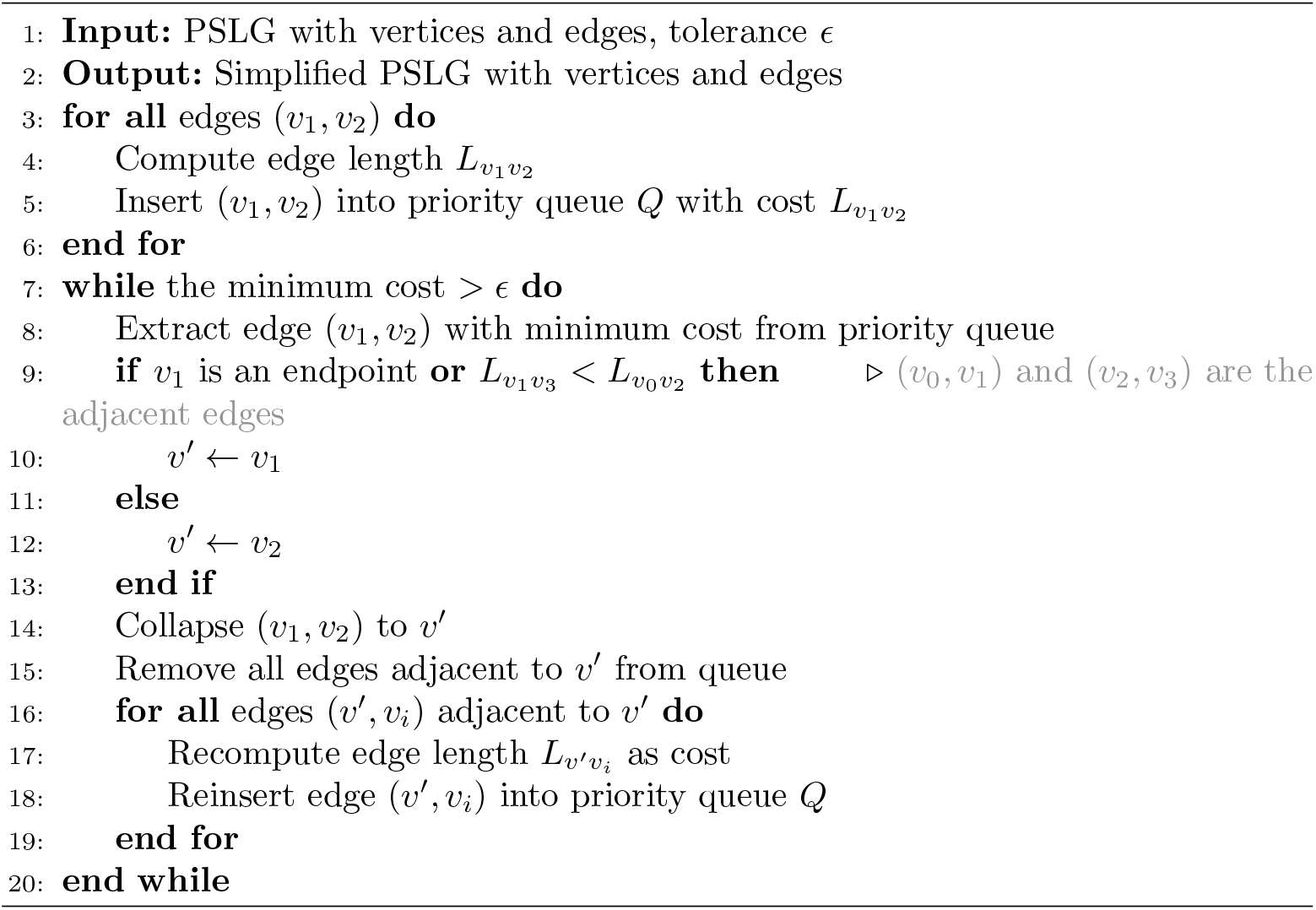

#### Lumped elements insertion

The lumped elements are modeled with the delta-gap method [62], which requires the lumped elements to be attached to edges that are aligned. We store the lumped element position as a parametric position *t* ∈ [0, 1) on the B-Spline. While converting B-splines into polylines, we first split the B-splines at these parametric positions, so that the lumped element position exists among the converted polyline vertices. We treat all the lumped element vertices as endpoints, so that they still exist after collapsing the small edges during strip generation. Finally, for each lumped element, we insert a line segment (the “delta gap”) that connects the two endpoints of the lumped element and attaches it to the strip PSLG. We also enforce that each lumped element is attached to a collection of edges forming a straight line, which is a requirement of MARIE.

#### Triangulation

We compute the Delaunay triangulation of the PSLG using the software Triangle [63]. During triangulation, we also save all the edge correspondences for the lumped elements by adding edge masks.

#### Mapping to 3D

Once the coils are discretized in 2D, the vertex positions of the triangulation are mapped to 3D using Φ(u). However, while coils may overlap in the 2D representation, they must not touch in 3D. We ensure this by adjusting our mapping function Φ(**u**) for each coil. The first coil *C*_0_ is mapped without any modifications, i.e., Φ_0_(**u**) = Φ(**u**). For every new coil we ensure that it does not intersect with the other coils already placed on the substrate. We achieve this by applying a displacement to the mapping,

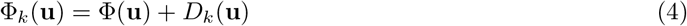

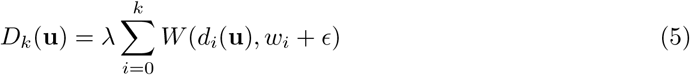

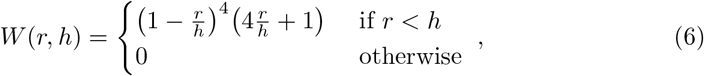

where *d*_*i*_(u) is the distance to the *i*^*th*^ coil, and *W* (*r, h*) is the Wendland function [64]. The parameters *λ* and *ϵ* control the height and the steepness of the displacement. The Wendland function has compact support and therefore influences the coil geometry only locally, creating bridges on the coil *C*_*k*_ (Figure 7).

**Fig. 7:**
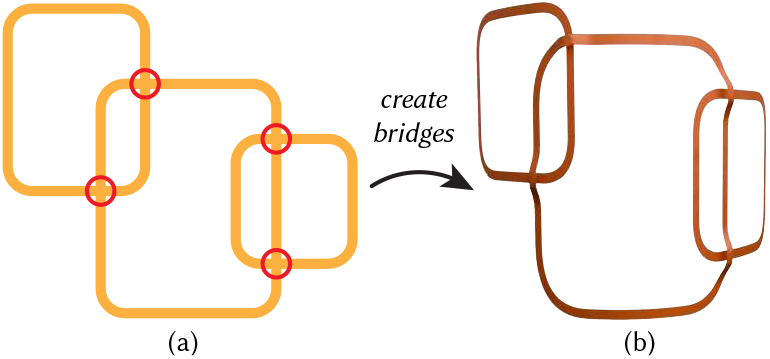
Coils can overlap in the 2D representation (a), but they must not intersect in the 3D model (b). We ensure this by embedding a displacement mapping that automatically creates bridges at positions where two coils cross each other.

## 5 Simulation

### 5.1 Background

Integral equation (IE) techniques offer distinct advantages for electromagnetic (EM) modeling in MRI applications. Unlike finite-difference time-domain and finite element methods, they are inherently free from grid dispersion artifacts [65, 66]. This is because IE formulations rely on Green’s functions, which serve as exact propagators of EM fields from sources to observation points. Furthermore, for time-harmonic (single-frequency) simulations, like MRI simulations, IE-based solvers give rise to dense matrices with properties such as symmetry, low-rank, hidden low-rank, or smooth spectral decay, which can be leveraged for fast and memory-efficient solutions using numerical linear algebra algorithms [67].

#### 5.1.1 Definitions

All EM simulations operate in periodic steady-state, at frequency *f* with units Hz. We define the angular frequency as *ω* = 2*π f* with units^rad^/_s_. In MRI, the frequency of operation is given by 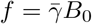, where 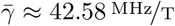 is the gyromagnetic ratio of ^1^H hydrogen atoms in water and B_0_ is the static magnetic field strength of the scanner. Given real-valued relative permittivity *ϵ*_R_, electrical conductivity σ, absolute permittivity in vacuum *ϵ*_0_, and imaginary unit *i*, we define the complex-valued relative permittivity as:

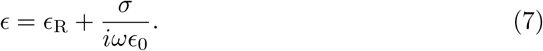

In the IE formulation, we also use the complex-valued electric susceptibility *χ* = *ϵ* − 1. In EM modeling, *ϵ, χ, ϵ*_R_, and σ are scalar fields, which we denote as 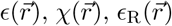, and 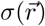, respectively. We discretize these quantities and the EM fields onto a uniform 3D grid of *N*_*v*_ voxels, and we denote them with a subscripted index to refer to their value at the voxel with the same index.

To describe coil tuning and decoupling, we extensively use the Schur complement, which is defined as follows: Given a block matrix,

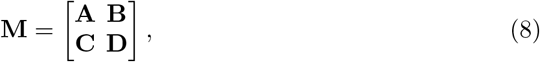

the Schur complement of block **A** in **M** is denoted as **M***/***A** and defined as:

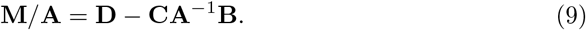

Similarly, the Schur complement of block **D** in **M** is defined as:

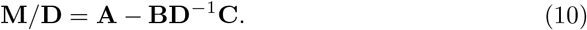

#### 5.1.2 Volume Integral Equation (VIE) solver

The Volume Integral Equation (VIE) formulation that we use involves calculating the volumetric current density 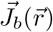 in the body subject to an incident electric field excitation at a single frequency *f*. The body is characterized by a distribution of relative permittivity 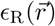 and electrical conductivity 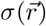, which we group into a distribution of complex relative permittivities 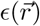.

To discretize the system, we can employ piecewise constant (PWC) basis functions and PWC testing functions, which results in a Galerkin discretization. We first consider a basis function centered at the origin *f*_0_ : ℝ^3^ → {0, 1} in a grid of voxel dimensions Δ_*x*_ × Δ_*y*_ × Δ_*z*_. We define *f*_0_ evaluated at point 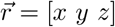 as:

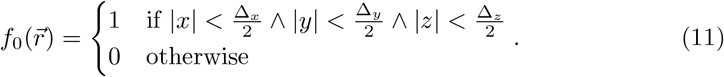

We then define the basis function *f*_*i*_ : ℝ^3^ → {0, 1}, at voxel *i* with center 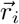:

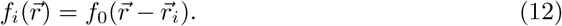

Note that each Cartesian component of the current density at a voxel has its own basis function, which means that the total number of PWC basis functions needed to discretize the current density is equal to 3*N*_*v*_.

We lump all possible body current contributions into density 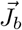 and consider this density to consist of equivalent current distributions in free space, for simplicity. The discretized system is given by:

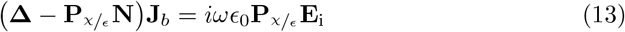

The term **E**_i_ refers to the discretized incident electric field distribution. The operator **P**_*χ*_/_*ϵ*_ multiplies each Cartesian component at voxel *i* by the corresponding ratio *χ*_*i*_/*ϵ*_*i*_. The operator **Δ** refers to the self-testing term or Gram matrix and for PWC basis functions is equal to Δ_*x*_Δ_*y*_Δ_*z*_**I**, where **I** is the identity matrix of appropriate dimensions.

With as few as 30 × 30 × 30 voxels for a 5 mm discretization, the full system would occupy approximately 100 GB of RAM when using double-precision arithmetic. To address this, we exploit the translation invariance of the Green’s function underlying the problem, and diagonalize the block Toeplitz operator **N** using the 3D Fast Fourier Transform (FFT), lowering the memory footprint from 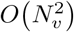 to *O*(*N*_*v*_). To invert the system, we can employ an iterative solver for non-symmetric systems, such as the Generalized Minimal Residual (GMRES) method.

#### 5.1.3 Surface Integral Equation (SIE) solver

The Surface Integral Equation formulation that we employ discretizes the single-frequency Electric Field Integral Equation (EFIE) using Rao-Wilton-Glisson (RWG) basis functions and RWG testing functions, resulting in a Galerkin discretization of the underlying system. Each RWG basis function is defined over a different pair of edge-adjacent triangles in the corresponding mesh.

Given a vector of basis coefficients **J**_*c*_ ∈ ℂ^*N*^, where *N* is the number of triangles in the mesh that discretizes the conductors, and a vector of applied voltages 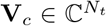 at the terminals, where *N*_*t*_ is the number of terminals (e.g., one port for each element of a coil array), the SIE system is given by:

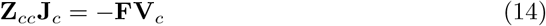

The complex symmetric matrix **Z**_*cc*_ ∈ ℂ^*N×N*^ is the Galerkin discretized system, and the operator 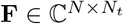 maps voltages to electric fields at the terminals. Assembling the **Z**_*cc*_ system involves computing *O*(*N*) singular and *O N*^2^ non-singular integrals. We adapt 1D Gauss-Legendre quadrature rules to compute these integrals, using a simple nesting scheme to approximate integrals over triangles. We compute the singular integrals using case-specific coordinate system transformations that remove the singularity from the integrand, at the expense of heavy use of trigonometric and inverse trigonometric functions.

Because the system **Z**_*cc*_ is complex symmetric and not Hermitian, we use direct inversion instead of conjugate gradients to invert the system in (14). Note that our SIE system results from the discretization of a Fredholm equation of the First Kind, which yields eigenvalues clustering at 0, further compromising the speed and applicability of iterative solving methods such as the conjugate gradients.

#### 5.1.4 Magnetic Resonance Integral Equation (MARIE) suite

In the Magnetic Resonance Integral Equation (MARIE) suite [68], SIE and VIE are jointly solved to compute the EM fields generated by RF coils within a dielectric sample (e.g., a numerical body model). The SIE models the coils by discretizing their geometry with triangular surface meshes and representing surface currents using RWG basis functions [69]. The VIE models the EM phenomena within the body, discretized on a uniform voxel grid, with polarization currents approximated via PWC basis functions [70]. The combined volume-surface integral equation (VSIE) framework exploits the multilevel Toeplitz structure of the Green’s function operators that map volumetric currents to fields, enabling fast matrix-vector multiplications through FFT [71].

We combine the SIE system with the VIE system by calculating the body-coil coupling operator 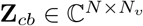, which maps from body current PWC basis coefficients **J**_*b*_ to an incident electric field for each RWG basis function. Similarly, we calculate the operator 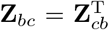 that maps RWG currents to incident electric field at each voxel. We introduce the term Z_*bb*_:

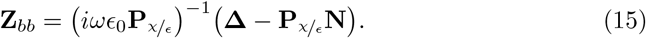

Finally, the coupled VSIE system is given by:

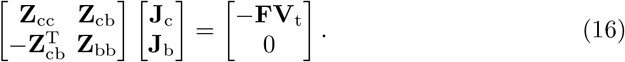

In the past decade, a series of algorithms have been implemented to further improve MARIE’s accuracy and efficiency. In terms of accuracy, it was shown that piecewise linear (PWL) basis functions yield more accurate simulations than PWC basis functions [72]. MARIE’s solution time can considerably increase for simulations involving fine voxel resolutions. To address this, since the discretized Green’s function tensors of the VIE sub-problem exhibit low multilinear ranks, it was shown that they can be compressed by at least two thousand times using the Tucker model, without sacrificing numerical precision [73]. As a result, the compressed operators can more easily fit in the limited memory of graphical processing units, leading to faster simulations. It was later shown that also the full VSIE system [74] can be compressed, using the precorrected FFT (pFFT) [75, 76] for cases when the coils are close to the body model, tensor train (TT) [77] for cases when the coils (or shields) are far from the body. A hybrid formulation of the VSIE that combines pFFT, cross-TT, and the adaptive cross approximation can also be used [78]. Finally, the SIE formulation has been adapted to explicitly model the coils’ lumped elements in the simulation [62] rather than considering each lumped element a separate port [37], which can considerably increase the number of required MARIE solves.

These advances have enabled a broader class of applications that involve iterative evaluations of MARIE or its constituent modules. In particular, the increased speed and accuracy of MARIE’s solver have allowed for the efficient use of body model-specific EM field bases to compute theoretical ultimate performance limits and the associated ideal current patterns (ICP) [11]. Tissue-dependent EM bases can be combined with the discrete empirical interpolation method [79] to construct the Magnetic Resonance Green Functions (MRGF) [80], which is a reduced-order model of MARIE’s VIE and VSIE operators.

#### 5.1.5 The Magnetic Resonance Green Function (MRGF)

EM simulation tools typically require an initial preprocessing step to assemble the necessary geometrical matrices involved in the simulation [80]. This preprocessing step can become computationally intensive, particularly for multi-element coil arrays discretized with thousands of triangles. For coil design optimization, where the coil geometry changes in each iteration, this pre-processing step becomes a significant bottleneck. The MRGF method addresses this high computational cost by allowing the incident fields of arbitrary RF coils to be expressed as linear combinations of precomputed fields of an EM basis [80]. This allows considerably faster predictions of the resulting electromagnetic fields in the body with less than 1% average accuracy loss when compared with the full MARIE solver [80]. Since MRGF transforms MARIE into a powerful framework for inverse problems, it was employed to implement the RF coil design optimization framework in this work.

Next, we briefly describe the steps associated with the MRGF method. First, one generates a basis of incident electric fields for a given body model by computing the fields generated by current sources external to the body. These current sources may consist of either a cloud of voxelized volumetric currents surrounding the body or of RWG surface currents whose support is a closed surface enclosing the body model [81]. One assembles the basis by exciting with one current element at a time and storing the vectorized incident field as a column of the incident field matrix 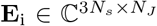, where *N*_*s*_ is the number of non-air voxels in the grid and *N*_*J*_ is the number of source current elements. Next, one applies the singular value decomposition (SVD) to **E**_i_, yielding the following factorization.

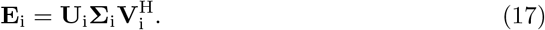

The orthonormal columns of the **U**_i_ matrix contain the dominant incident field modes, which one would then truncate up to a desired tolerance in the matrix *L*_2_ norm sense, or equivalently one would keep all vectors whose singular values *σ*_*k*_ satisfy *σ*_*k*_ *> σ* _1_*ε*, where *σ*_1_ is the largest singular value and *ε* is the prescribed tolerance.

After the calculation of the incident field basis, one applies the Discrete Empirical Interpolation Method (DEIM) [79] to select a subset of voxels for which to calculate the incident fields, which can then be interpolated to derive the incident fields for all tissue voxels using the operator **X** that DEIM provides. After DEIM, one solves the VIE system **Z**_bb_ for all incident fields in **U**_i_, yielding a set of volumetric currents stored in matrix **M**. One then forms the matrix **M**_m_:

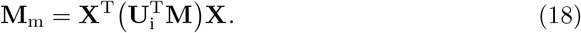

When simulating a coil geometry, one would calculate the matrix **Z**_dc_, which is equal to the incident electric field at the DEIM interpolation points over all of the RWG coil currents. The product 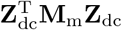 is a close approximation of the term 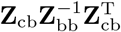 of the full VSIE system. One then applies the SVD to **M**_m_, resulting in the approximate factorization 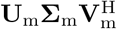, which one would then truncate to achieve a specified tolerance, as was done with the incident fields in **U**_i_, concluding the MRGF assembly process.

The MRGF only needs to be assembled one time per body model, and can be used to simulate any coil configuration constrained to surfaces that lie or extend beyond the substrate where the current sources used to generate the basis were defined. In summary, using the MRGF method, we can precompute the inverse of the VIE operators and reduced the number of points needed to compute the electric and magnetic field over all voxels, drastically accelerating the solution of (16), which we invert directly following the procedure outlined in [80]. With this precomputed solution, we then calculate the total electric and magnetic fields, which are required to determine the SNR during coil design optimization.

### 5.2 Simulation improvements

The previous versions of MARIE [37, 76] assembled the SIE operators using fixed quadrature orders that were too low for precise computation of the coil-to-coil interaction matrix **Z**_cc_. A lack of precision in these operators means that gradient calculations via finite differences will be too inaccurate for optimization, unless the step size is increased substantially. This code was additionally unoptimized and implemented in the MATLAB scripting language, resulting in assembly times for **Z**_cc_ that would render the optimization intractable. Additionally, the previous version lacked algorithms for tuning, matching, and decoupling, which are essential components of any electromagnetic simulations involving coils and antennas. In this work, we developed optimized routines for the assembly of **Z**_cc_ that are not only faster but also considerably more precise. We also developed algorithms for tuning and decoupling of coil arrays that allow us to simulate the coil-to-coil interactions more realistically.

#### 5.2.1 Improving the precision of the SIE operator assembly

The SIE assembly process for computing coil-to-coil interactions **Z**_cc_ involves numerically integrating complex-valued singular functions for four different scenarios: vertex adjacent pairs, edge adjacent pairs, overlapping or self-term pairs, and non-singular pairs. Since the non-singular pairs are analytic over the 4D integration domains, we apply simple Gauss-Legendre quadrature to integrate these terms. However, for the other three cases, we apply coordinate transformations that then eliminate these singularities at the expense of greater numerical complexity in the integrands. We start by considering the real part of an integrand used in the vertex-adjacent integrals,

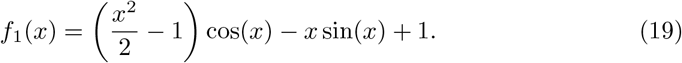

When evaluating these integrands, *x* is equal to *k*_0_*r*, where *k*_0_ is the wavenumber in free space and *r* is the distance between observation and source quadrature points. Such a function, when evaluated for small arguments as defined, suffers from catastrophic loss of numerical precision. We can understand why by expressing this function in terms of its Taylor series about *x* = 0, which has an infinite radius of convergence in *x*:

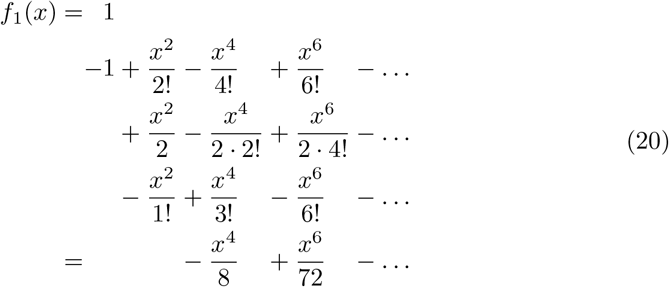

As is evident, the constant and quadratic terms in the Taylor series cancel out. For small arguments, the quartic terms are considerably smaller than the quadratic terms.

When these terms cancel out, most of the precision in the floating point representation is lost, resulting in a catastrophic loss of numerical precision. Figure 9 shows the relative error in the direct evaluation of *f*_1_(*x*) when using double-precision floating point numbers with respect to the direct evaluation when using 512-bit precision floating point numbers, as well as the relative error of a Taylor Series approximation with 12 non-zero coefficients (polynomial order 26). Figure 8 shows the convergence of the relative error in **Z**_cc_ as a function of Gauss-Legendre quadrature order, for each different adjacency type. For the quadrature orders in the original implementation, the relative error in **Z**_cc_ was on the order of 10^*−*4^, which is insufficient for coil optimization. Additionally, even when the quadrature order was increased to the maximum possible order, the relative error in **Z**_cc_ plateaued at approximately 10^*−*8^, which was still significant, potentially impacting the estimation of gradients and the line search procedure when optimizing coil array geometries.

**Fig. 8:**
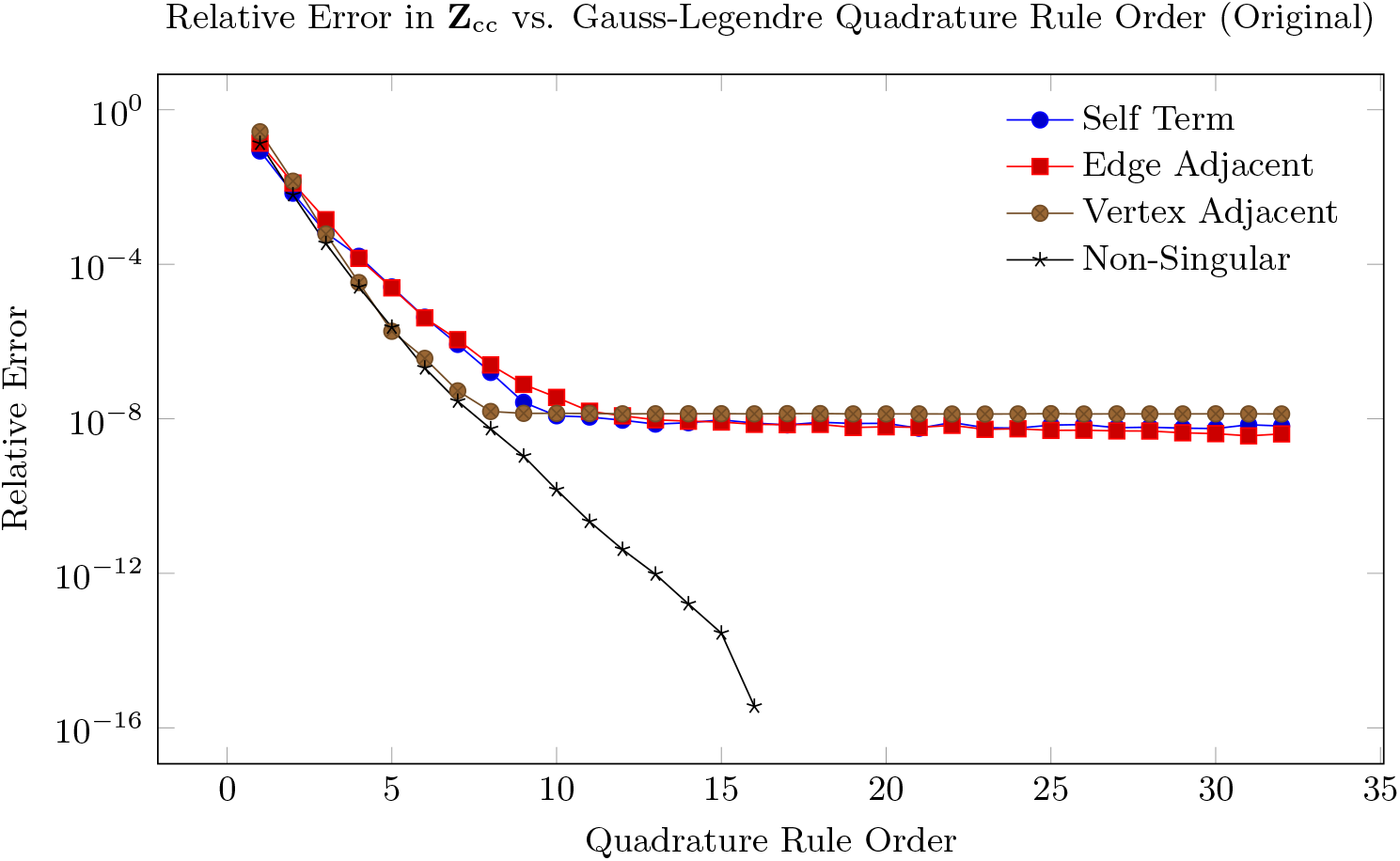
Convergence of error in SIE integrals as a function of Gauss-Legendre quadrature order, when assembling Z_cc_ using the original implementation.

**Fig. 9:**
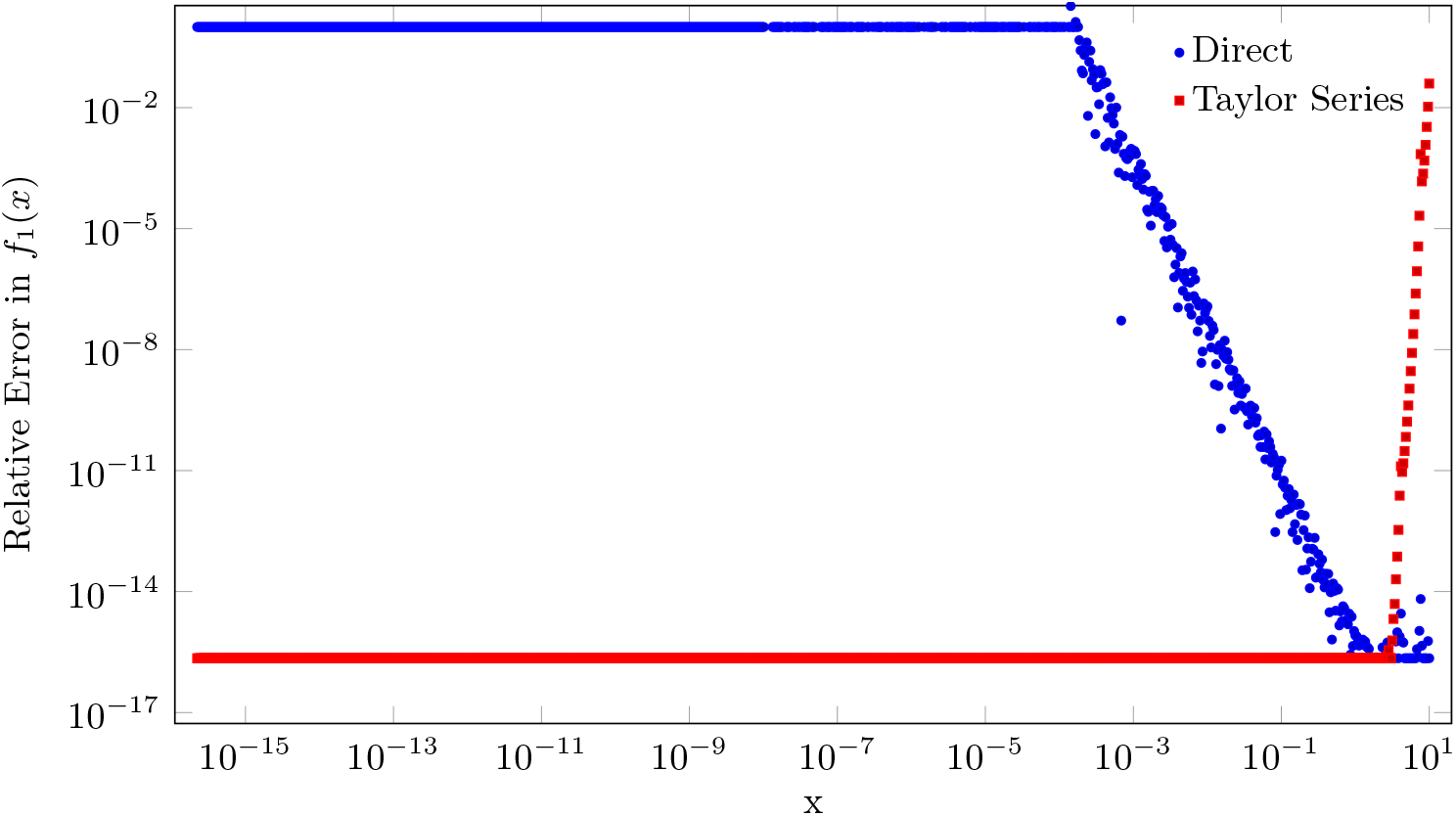
Evaluation of *f*_1_(*x*) using direct evaluation (blue) and Taylor Series approximation (red). The precision of direct evaluation suffers from a total loss of precision for arguments of magnitude 10^−3^ or smaller. The threshold is set to the point in the graph where the two sets of data meet (*x* ≈ 3).

We addressed this loss of numerical precision by dividing the evaluation of each singular integrand into two cases depending on whether the absolute value of the argument is smaller or larger than a threshold. The threshold depends on the approximated function but is usually either |*x*| = 1 or |*x*| = 3. For this work, it was determined empirically for every integrand by comparing the evaluation in double-precision with the evaluation in either 512-bit floating point representations for logarithmically spaced samples. When the input exceeds the threshold, we evaluate the integrands using their functional forms, as in (19). When the input is smaller than the threshold, we use the Taylor series of the integrand, evaluated using Horner’s method for polynomial evaluation, because direct polynomial evaluation can also suffer from loss of precision. We summarize all of the replaced integrands in the SIE assembly and their Taylor series approximations in Table 1. We include the leading term of each Taylor series about 0, as the order of the leading term correlates with the severity of the loss of numerical precision. We also subdivided the non-singular interactions into four categories, based on the ratio of the distance between the triangle pairs to the free-space wavelength *λ* =

**Table 1:** Singular integrals needed for the SIE assembly and associated Taylor expansion when the input *x* is lower than the specified threshold.

| Function | Leading Term | Threshold |
| --- | --- | --- |
| $\cos(x) - 1 + \frac{x^2}{2}$ | $+\frac{x^4}{4!}$ | 1 |
| $\cos(x) - 1 + \frac{x^2}{2} - \frac{x^4}{24}$ | $-\frac{x^6}{6!}$ | 1 |
| $x - \sin(x)$ | $+\frac{x^3}{3!}$ | 1 |
| $x - \frac{x^3}{3!} - \sin(x)$ | $-\frac{x^5}{5!}$ | 3 |
| $\cos(x) + \frac{x}{2} \sin(x) - 1$ | $-\frac{x^4}{4!}$ | 3 |
| $\cos(x) + \frac{x}{2} \sin(x) - 1 + \frac{x^4}{4!}$ | $+\frac{2x^6}{6!}$ | 3 |
| $\frac{x}{2}(\cos(x) + 1) - \sin(x) + \frac{x^3}{12}$ | $+\frac{x^4}{80}$ | 3 |
| $\frac{x}{2}(\cos(x) + 1) - \sin(x)$ | $-\frac{x^3}{12}$ | 1 |
| $\left(1 - \frac{x^2}{8}\right) \cos(x) + \frac{5x}{8} \sin(x) - 1$ | $-\frac{x^6}{6!}$ | 3 |
| $\frac{3x}{8} + \frac{x^3}{48} + \frac{5x}{8} \cos(x) - \left(1 - \frac{x^2}{8}\right) \sin(x)$ | $-\frac{x^5}{320}$ | 3 |
| $\left(3 - \frac{x^2}{2}\right) \cos(x) + 2x \sin(x) - 3 - \frac{x^4}{4!}$ | $-\frac{x^6}{5!}$ | 3 |
| $2x \cos(x) + \left(\frac{x^2}{2} - 3\right) \sin(x) + x$ | $-\frac{3x^5}{5!}$ | 3 |
| $\left(1 - \frac{x^2}{8}\right) \cos(x) + \frac{3x}{4} \sin(x) - 1 - \frac{x^2}{8} + \frac{x^4}{48}$ | $-\frac{x^6}{4 \cdot 6!}$ | 3 |
| $\left(\frac{x^2}{8} - 1\right) \sin(x) + \frac{3x}{4} \cos(x) + \frac{x}{4} + \frac{x^3}{12}$ | $+\frac{x^5}{4 \cdot 5!}$ | 3 |
| $\frac{x}{2} \cos(x) + \left(\frac{x^2}{8} - 1\right) \sin(x) + \frac{x}{2} - \frac{x^3}{4!}$ | $-\frac{x^5}{5!}$ | 3 |
| $\cos(x) - 1 + \frac{x}{4} \sin(x) + \frac{x^2}{4}$ | $+\frac{x^6}{2 \cdot 6!}$ | 3 |
| $\frac{x}{4} \cos(x) - \sin(x) + \frac{3x}{4} - \frac{x^3}{4!}$ | $+\frac{x^5}{4 \cdot 5!}$ | 3 |
| $\cos(x) + x \sin(x) - 1$ | $+\frac{x^2}{2}$ | 1.6818 |
| $\cos(x) + x \sin(x) - 1 - \frac{x^2}{2}$ | $-\frac{x^4}{8}$ | 3 |
| $x \cos(x) - \sin(x)$ | $-\frac{x^3}{3}$ | 3 |
| $\left(1 - \frac{x^2}{2}\right) \cos(x) + x \sin(x) - 1$ | $+\frac{x^4}{8}$ | 3 |
| $x \cos(x) + \left(\frac{x^2}{2} - 1\right) \sin(x)$ | $+\frac{x^3}{6}$ | 1.6818 |
| $\left(1 - \frac{x^2}{2}\right) \cos(x) + \left(x - \frac{x^3}{6}\right) \sin(x) - 1$ | $-\frac{x^4}{4!}$ | 1.6818 |
| $\left(x - \frac{x^3}{6}\right) \cos(x) + \left(\frac{x^2}{2} - 1\right) \sin(x)$ | $+\frac{4x^5}{5!}$ | 3 |
| $\cos(x) + \frac{x}{3} \sin(x) + \frac{x^2}{6} - 1$ | $-\frac{x^4}{3 \cdot 4!}$ | 3 |
| $\frac{x}{3} \cos(x) - \sin(x) + \frac{2x}{3}$ | $+\frac{2x^5}{3 \cdot 5!}$ | 3 |
| $\left(1 - \frac{x^2}{6}\right) \cos(x) + \frac{2x}{3} \sin(x) - 1$ | $+\frac{x^4}{3 \cdot 4!}$ | 2 |
| $\left(\frac{x^2}{6} - 1\right) \sin(x) + \frac{2x}{3} \cos(x) + \frac{x}{3}$ | $-\frac{x^5}{5!}$ | 3 |
| $\left(\frac{x^2}{2} - 1\right) \cos(x) - x \sin(x) + 1$ | $-\frac{x^4}{8}$ | 3 |
| $\left(1 - \frac{x^2}{2}\right) \sin(x) - x \cos(x)$ | $-\frac{x^3}{6}$ | 3 |
| $\left(\frac{x^2}{2} - 1\right) \cos(x) - \left(\frac{x^3}{6} - x\right) \sin(x) + 1$ | $+\frac{x^4}{4!}$ | 1.6818 |
| $\left(1 - \frac{x^2}{2}\right) \sin(x) + \left(\frac{x^3}{6} - x\right) \cos(x)$ | $-\frac{x^5}{30}$ | 3 |
| $\left(-\frac{x^4}{24} + \frac{x^2}{2} - 1\right) \cos(x) + \left(\frac{x^3}{6} - x\right) \sin(x) + 1$ | $+\frac{x^6}{144}$ | 3 |
| $\left(+\frac{x^4}{24} - \frac{x^2}{2} + 1\right) \sin(x) + \left(\frac{x^3}{6} - x\right) \cos(x)$ | $+\frac{x^5}{5!}$ | 3 |

Fig. 10 demonstrates the same convergence analysis as in Fig. 8 but with the proposed implementation.

**Fig. 10:**
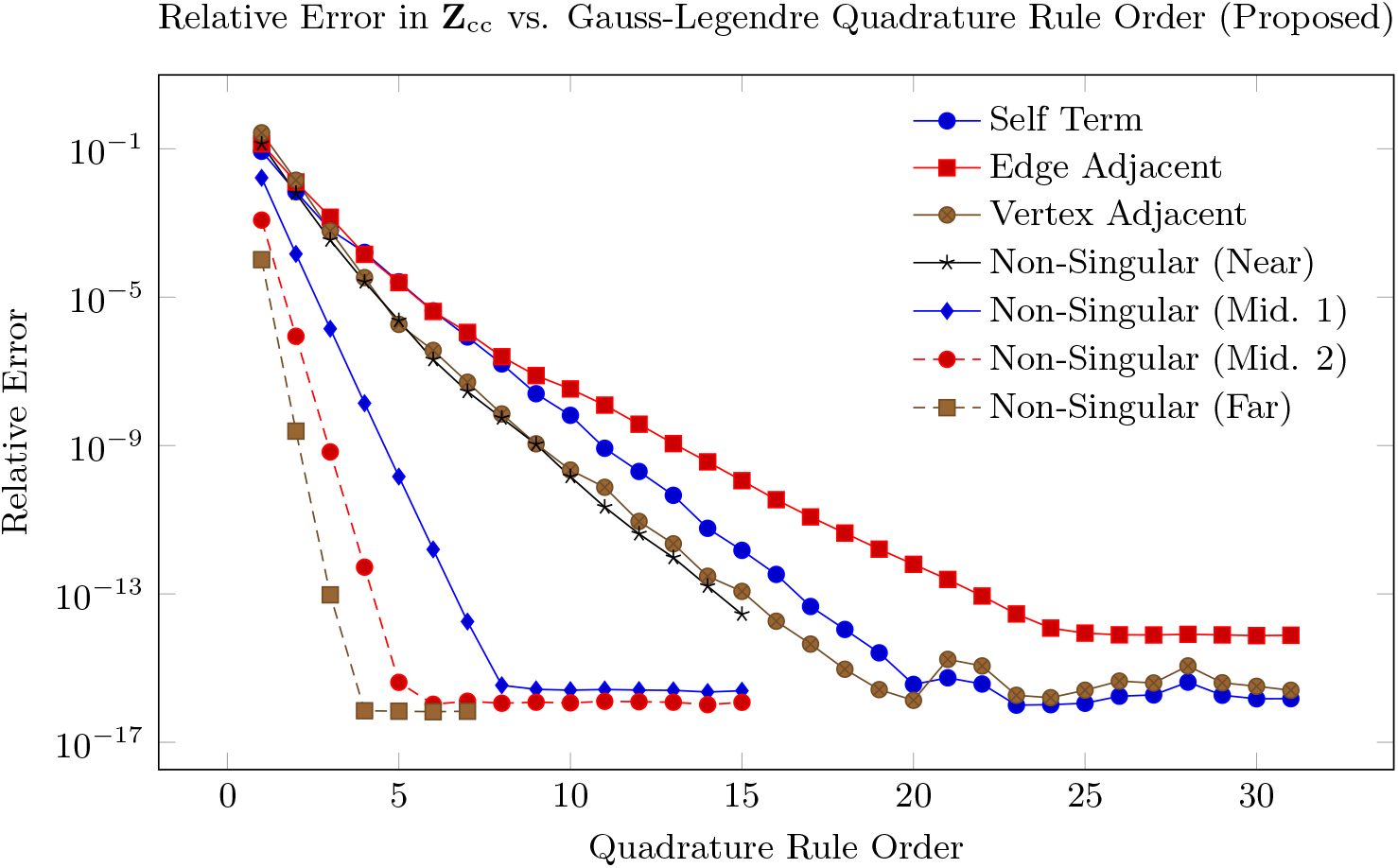
Convergence of error in SIE integrals as a function of Gauss-Legendre quadrature order, when assembling **Z**_cc_ using the proposed implementation.

**Fig. 11:**
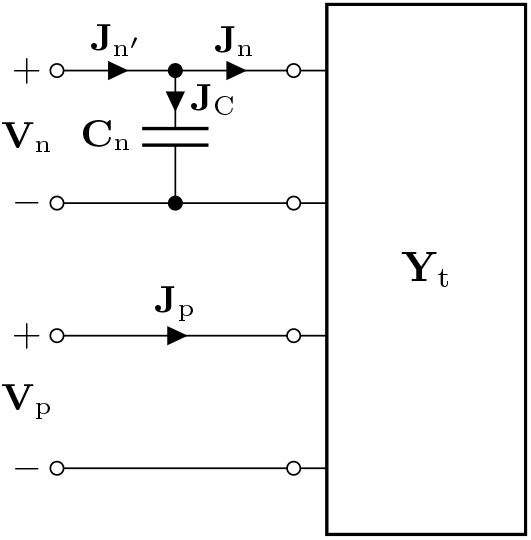
Circuit schematic demonstrating the tuning approach.

#### 5.2.2 Accelerating the assembly of SIE Operators

The original SIE assembly implementation was written in the MATLAB scripting language and was unoptimized. As we saw in the previous section, many of the integrands in the calculation of singular integrals also suffered from catastrophic loss of numerical precision, necessitating custom implementations of these functions for small arguments that are also fast to compute. We addressed this requirement and the need for optimized code by rewriting the SIE assembly code in C++. We introduced a number of optimizations meant to both decrease assembly times and to lighten the load on CPU caches.

Each singular integrand type uses a number of trigonometric expansions that are independent of the dimensions of the pairs of triangles, and depend only on the quadrature order. We calculated these geometry-independent quantities at compile time using the C++11 standard’s `constexpr` variable and function modifier, which forces evaluation at compile time. Additionally, the original implementation kept a sorted list of non-singular interaction pairs over the *N*_*t*_ triangles of the mesh that grows as 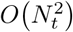, as well as sorted lists of vertex-adjacent, edge-adjacent, and self term pairs that grow as *O*(*N*_*t*_). The self term pairs consist of pairs {(*i, i*) for *i* ∈ {1, 2, …, *N*_*t*_}}, which can be inferred automatically during assembly. The non-singular pairs consist of all pairs that are not vertex-adjacent, edge-adjacent, or self terms. This allows us to discard entirely the list of non-singular pairs, which we replace with a loop that iterates over all possible pairs, using a binary search algorithm over the entries in the vertex-adjacent and edge-adjacent lists to determine whether to carry out the computation. The binary search introduces a negligible *O*(log(*N*_*t*_)) term during assembly, but because the input data needed to assemble the operator grows as *O*(*N*_*t*_) instead of 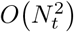, the assembly is typically considerably faster due to improved cache coherence.

### 5.3 Admittance and impedance parameters over terminals

When we apply voltages **V**_t_ to the coil array, we obtain a corresponding set of currents **J**_t_, with positive currents flowing into the corresponding positive side of each terminal. The linear operator describing this transfer function is called the admittance parameter matrix, which we denote as **Y**_t_. The voltage-current relationship can thus be summarized as

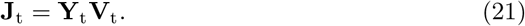

In order to calculate the admittance parameters **Y**_t_, we use the Schur complement of block **Z**_bb_ in (16) to obtain the loaded coil-to-coil interaction matrix **Z**_cbc_,

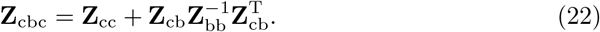

We can thus express the solution for J_t_ in (16) as

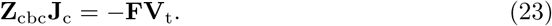

Finally, we use the identity **J**_t_ = −**F**^T^**J**_c_ to obtain the following relationship between applied terminal voltages **V**_t_ and induced terminal currents **J**_t_. The minus sign stems from **F**^T^**J**_c_ resulting in the current flowing out of the positive side of the terminal,

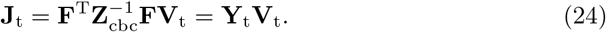

The operator acting on **V**_t_ is precisely the admittance parameter matrix of the system.

#### 5.3.1 Accelerating SVIE using the MRGF

As discussed in Sec. 5.1.5, we used the MRGF method to drastically accelerate the computation of **Z**_cbc_ in (22). More specifically, for each coil geometry, we calculated the incident electric field over the DEIM interpolation points **Z**_dc_ and computed the product with

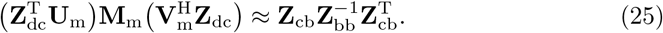

The right-hand side is precisely the SVIE term appearing in (22), which we replace with the left-hand side to calculate the SVIE term much more efficiently with the precomputed VIE inverse operator **M**_m_. In this work, we required fewer than 5, 000 interpolation points when assembling the MRGF operators for all numerical experiments at 5 mm isotropic resolution.

### 5.4 Adding tuning capacitors to coils

The terminals can be subdivided into two categories: non-port terminals and port terminals. We denote each type using subscripts *n* and *p*, respectively. As such, we can subdivide the terminal voltages, terminal currents, and admittance (or impedance) parameters using this convention, allowing us to rewrite (24) as follows:

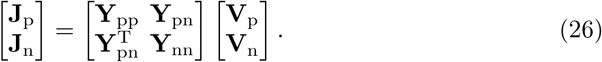

In order to tune the coil array, we attach capacitors C_n_ at the non-port terminals in parallel. Consequently, when we express the admittance parameters of (26) with a new non-port terminal current 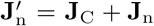 flowing into the parallel combination, with **J**_C_ corresponding to the vector of currents flowing into the capacitors. The Voltage-Current (V-I) relationships of the capacitors are

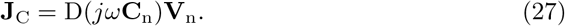

We can thus write the current-voltage (I-V) relations with respect to J_n_′,

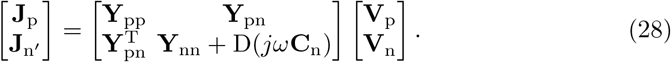

We introduce **Y**_nn_′ = **Y**_nn_ + D(*j ω* **C**_n_) and obtain the loaded impedance parameters by applying the Schur complements to invert the loaded admittance matrix over the terminals,

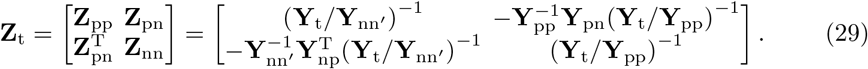

This square matrix dictates the Voltage-Current (V-I) relationships of the loaded system. However, because only the capacitors at the non-port terminals load the coil array, no current will flow into the primed terminals, meaning that **J**_n_′ = 0, which allows us to ignore the second block column,

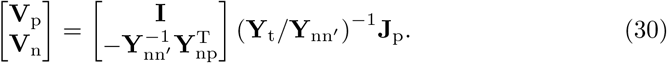

The top block dictates the Voltage-Current relationships over the ports and is equal to the impedance parameter matrix over the ports **Z**_p_. The admittance parameters over the ports, which we use in the following section when tuning, are simply the inverse of **Z**_p_,

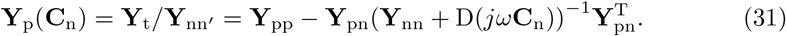

While not relevant when tuning, the lower block contains the transfer function from the port currents **J**_p_ to the voltages at the non-port terminals, which we will use when discussing our decoupling strategy.

#### 5.4.1 Tuning a coil array

We apply Lorentz reciprocity to our problem and argue that decoupling preamplifiers behave like current sources in parallel with source resistances under reciprocity. We set these resistances to ∞ and refer to this process as *ideal decoupling*. Note that this could be approximated in practice for receive arrays with preamplifier decoupling. As a result, when a port is driven, all of the other port currents are identically 0. Therefore, we only consider the self-interactions when tuning the coil array, i.e., we consider only the diagonal of loaded impedance parameters **Y**_p_ in (31). We tune the coil array by solving the following optimization problem:

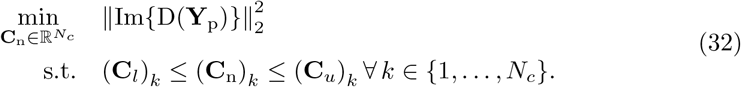

The vectors C_*l*_ and C_*u*_ denote lower and upper bounds, respectively, for the tuning capacitors. This optimization problem is highly non-convex and features sharp peaks near the optima with flat regions between these optima. When the cost function exceeds a pre-determined threshold, we solve this problem using a Particle Swarm optimization. Once that optimization completes, we refine the solution by solving the optimization problem using Newton’s Method.

#### 5.4.2 Decoupling a coil array

When we solve the coupled SVIE system, we simply invert the operator on the lefthand side of (23) while setting **V**_t_ to the identity matrix. This is equivalent to setting the voltage at a port to 1 V and the rest to 0 V using voltage sources, and repeating this for all ports. We denote this solution as **ĵ**_c_, which we further subdivide,

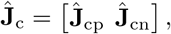

based on whether the voltage was applied at a port or a non-port terminal. We then perform tuning, resulting in a set of capacitors C_n_. Given these capacitors, we can obtain the voltage at each terminal using the transfer function in (30) that maps from port currents to terminal voltages. We then multiply our original solution **ĵ**_c_ by this transfer function to obtain the RWG basis coefficients corresponding to driving a tuned, decoupled coil using current sources,

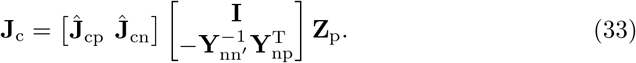

## 6 Optimization

We cast the optimization of an RF coil as a constrained optimization problem. We define an objective function *f* (*x*), which evaluates the coil’s absolute performance in an ROI, along with a constraint function *g*(*x*) that ensures that the coil geometry is valid (Section 3). Whenever a coil is considered invalid, *g*(*x*) is negative, e.g., for the case of a coil intersecting itself or other coils. Our coil shape optimization problem can therefore be written as:

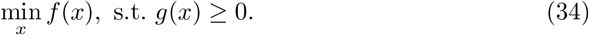

We convert this into an unconstrained optimization problem where the constraints are enforced by a barrier potential term *B*_*g*_(*x*) that increases to infinity if *g*(*x*) approaches 0, and vanishes when *g*(*x*) *>* 0 is sufficiently large.

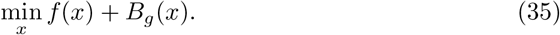

The conversion to the unconstrained problem using a potential has several advantages: First, while inequality constraints may introduce non-smoothness to the solution, replacing these with a smooth barrier in general leads to approximating smooth solutions of similar accuracy, as long as the underlying PDEs allow for this. Second, converting the problem to an unconstrained optimization problem allows one to use simpler and more reliable optimization algorithms. On the downside, barrier potentials introduce additional non-physical forces in configurations close to contact. However, if the extent of barrier potential is chosen to be sufficiently small, the overall solution error is dominated by the geometry discretization error.

We solve the optimization problem in (35) using L-BFGS [82]. The gradients for the barrier term are computed analytically. The gradients for the objective are estimated using finite differences. Although an analytic derivation of the gradient of *f* (*x*) is possible, one would need to compute multiple derivatives of multidimensional singular integrals appearing in the MRGF’s geometrical Green’s function operators [80]. For simplicity, this work employs the finite difference method. In future work, we will consider analytic gradients to make searching for a larger design parameter space computationally manageable. Finally, we use explicit line-search checks to ensure that the optimization remains within the feasible space *g*(*x*) *>* 0 (Section 6.2).

### 6.1 Objective function

Our goal is to optimize receive coils so that the associated SNR approaches the UISNR within a target region of interest (ROI). Once the electric (e) and magnetic (h) fields of each coil element are computed using an EM simulator (e.g., MARIE or MRGF described in 5), the optimal SNR at a position of interest r_0_ ∈ ROI can be computed as [5]:

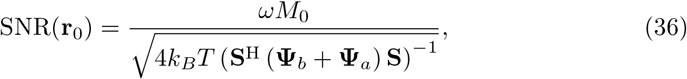

with *ω* the angular operating frequency, *M*_0_ the equilibrium magnetization, S ∈ ℂ^*n×p*^ the receive coil sensitivity (= h_x_ − ih_y_), *k*_*B*_ Boltzmann’s constant, *T* the temperature of the sample, *n* the number of coils in the array, and *p* the

The noise covariance matrix Ψ_*b*_ ∈ ℝ^*n×n*^ [83] accounts for intrinsic thermal losses due to the sample’s conductivity and its elements can be computed for each coil pair *n*_1_, *n*_2_ as:

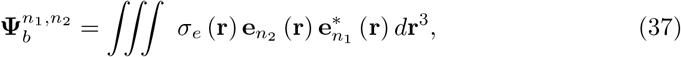

where the integral is computed over the entire sample and *σ* _*e*_ ∈ ℝ^*p×*1^ is the electric conductivity. Ψ_*a*_ ∈ ℝ^*n×n*^ is a diagonal matrix computed using the equivalent-power formulation presented in [84], which models the noise equivalent resistance associated with Joule heating of the coil conductors. The RF coils are modeled as non-perfect electric conductors by incorporating the finite surface resistivity of copper directly into the surface integral equation (SIE) matrix, following the formulation in [62], to account for ohmic losses in the conductors.

The UISNR is computed using Equation 36, without Ψ_*a*_, combining the elements of an EM basis as if they were coils in a hypothetical infinite array. One can construct a basis of EM fields following the Huygens–Fresnel principle [85] and the approach in [81]. This basis includes all possible field distributions within the sample generated by RF sources placed outside an enclosing surface. As the number of modes included in Equation 36 increases, the resulting SNR converges to the UISNR [86–88], which represents the theoretical maximum achievable SNR for the given sample, independent from coil geometry. The UISNR must be computed once as the benchmark of the optimization problem, while the SNR is updated in every iteration of the optimization for each new coil configuration.

For our problem, the objective function (Equation 35) quantifies the difference between the simulated SNR and the UISNR within a target ROI indicated by r,

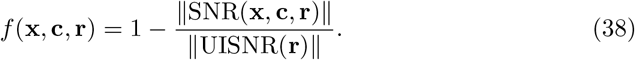

Here, x represents the geometric parameters of the coils, and c are the variable capacitors distributed around the coil conductors, which are optimized for coil tuning (see Section 5.4.1). Note that, depending on the sample and frequency of operation, the UISNR values could diverge at voxels near the surface of the sample due to numerical instability [89]. To ensure the validity of our results, the ROIs considered in this work do not include voxels located within 2.5 cm of the surface of the sample.

For a meshed coil, we define the constraint *g*(*x*) as

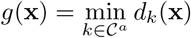

where 𝒞^*a*^ is the set of all possible pairs (*e, v*) of edges *e* and vertices *v* on the boundaries of coils, and *d*_*k*_ is the signed distance between a vertex and an edge in the pair *k*, where the sign is determined by the dot product with edge perpendicular pointing outside the coil.

Note that we do not compute *g*(*x*) explicitly; rather, we define a potential *B*_*g*_ that ensures that *g*(x) remains positive, and much less expensive to compute, as it uses only nearby pairs.

We constrain the objective function with a barrier potential term *B*_*g*_(*x*) that forces the coils to stay within the feasible domain, and prevent individual coils from self-intersecting (Section 3):

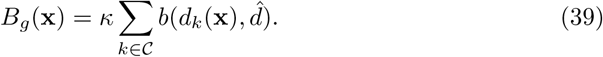

*B*_*g*_(*x*) measures the sum of these barriers over each contact pair *k* ∈ 𝒞 ⊂ 𝒞^*a*^, which are sufficiently close. The parameter *κ >* 0 controls the barrier stiffness, which primarily affects the optimization efficiency. Following [54], we construct a continuous barrier energy *b* that creates a localized repulsion force when primitives are closer than a distance 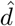, and that vanishes otherwise (see Figure 1).

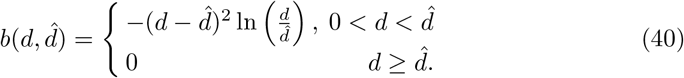

In the definition of *B*_*g*_, the set 𝒞 can be restricted to pairs which are closer than 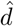. We use the logarithmic potential following [54]; this choice is far from unique, as long as the essential barrier properties are satisfied (infinite barrier at contact, zero value at a small distance to contact and smoothness w.r.t., geometric degrees of freedom). The logarithmic potential was demonstrated to work well in [54] and follow-up works, and is also commonly used in interior-point solvers for constrained optimization.

### 6.2 Optimization process and line search

The objective function (38) and the barrier potential term (39) are both smooth, allowing gradient-based optimization to be employed. The optimization algorithm consists of two steps. First, we optimize the size and location of the coils with only affine transformations by defining a scaling factor *s* and translation distances *t*_*u*_ and *t*_*v*_ for each coil. Second, we further optimize individual B-spline control point coordinates (*p*_*u*_, *p*_*v*_). The first step has 3 × *n* parameters, and the second step has 2 × *m* parameters to optimize, where *n* is the number of coils and *m* is the total number of control points.

In each step, we start with a grid search of the parameters, using *q* samples on each dimension, which requires *O*(*q*^*n*^) simulations. To reduce computational load, we use coordinate descent, i.e., we search for the optimal solution in one dimension at a time, and terminate when the objective can no longer decrease in any dimension. This reduces the complexity to *O*(*nq*) simulations.

After the grid search, we continue with gradient-based optimization with gradient descent or L-BFGS. We compute the gradient of the objective from (35) by adding ∇*f* (*x*), computed with finite differences, and the gradient of the barrier potential ∇*B*_*g*_(*x*). ∇*B*_*g*_(*x*) is derived with a chain rule by multiplying the analytic derivative D*B*_*g*_ of the barrier potential term [54] and the shape velocity D*v* defined on mesh vertices with respect to the optimization parameters:

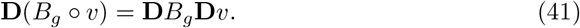

Next, we propose a continuous, intersection-aware line search for the optimization parameters. In each line search, we first apply an intersection-aware continuous collision detection (CCD) [54, 90] (StepSizeUpperBound in Algorithm 2) to conservatively compute the largest feasible step size along the gradient descent direction. We then apply a backtracking line search, bound by the largest feasible step size, to obtain an energy decrease. We terminate the optimization when the norm of gradient is smaller than *ϵ*_*d*_ = 10^*−*4^ or the line search fails to find an energy decrease with a maximum number of iterations *n*_*d*_ = 30 (Algorithm 2).

## 7 Methods

We first demonstrated the numerical issues that we addressed with the improved precision in the SIE assembly code (Section 5.2.1) by estimating gradients of the SNR cost function (Eq. (38)) of a simple parametrization of a single loop. We then conducted numerical experiments of increasing complexity to demonstrate the coil design optimization pipeline. We started by optimizing a single parameter – the horizontal placement of a single electric dipole (Section 7.3) – as we moved the target ROI. We then optimized the position and shape of three coils to maximize the average SNR within a spherical ROI at different locations inside a head model (Section 7.4). To evaluate our framework for different cost functions, we optimized four coils to maximize SNR on two decoupled ROIs (Section 7.5). Finally, we optimized a 12-coil head array for different ROIs in the brain (Section 7.6) and tested the generalization capacity of our optimization approach by evaluating the consistency of the results for a different head model. All computations were executed on a server running the Ubuntu 24.04.2

### Algorithm 2 Gradient-based Optimization. ComputeConstraintSet determines the set of pairs of segments for which the contact potential is not zero.

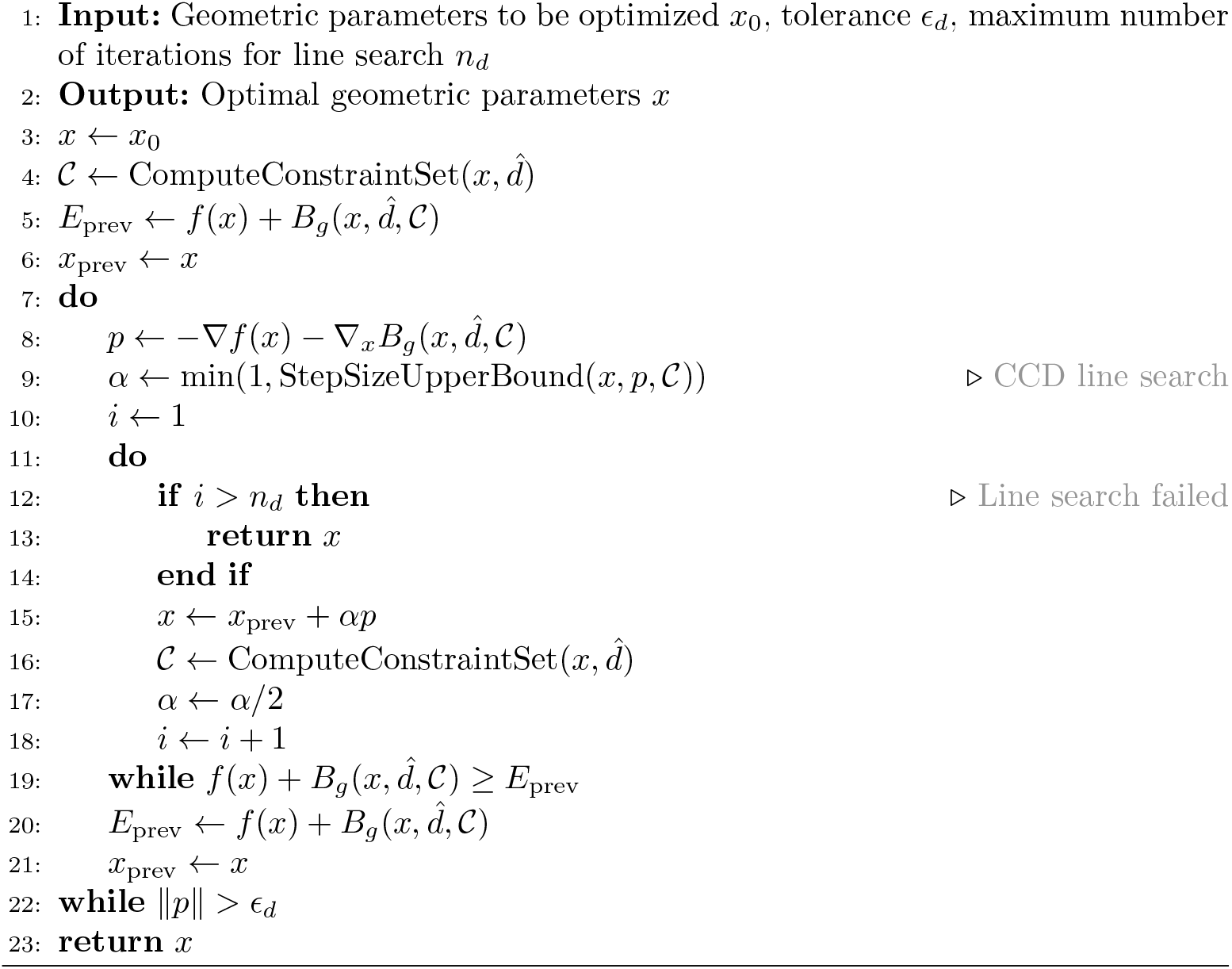

LTS operating system, equipped with an AMD Ryzen Threadripper PRO 3995WX CPU at 2.70 GHz, 64 cores, 2 threads per core and an NVIDIA GeForce RTX 3080 Ti GPU with 12 GB of memory.

### 7.1 Numerical samples

The experiments in Sections 7.3, 7.4, 7.5, 7.6 used the realistic Duke human model of the Virtual Family Population [91]. The computational domain enclosing the Duke’s head model was 18.5 × 23 × 22.5 cm^3^ and was discretized over a uniform grid of 5 mm^3^ voxel resolution, corresponding to 38 × 47 × 46 voxels. For one experiment in Section 7.6, we used the Ella human model of the Virtual Family with a computational domain of 17.5 × 21 × 24 cm^3^, which was discretized over a uniform grid of 5 mm^3^ voxel resolution, corresponding to 35 × 42 × 48 voxels.

### 7.2 Coil formers

To calculate the UISNR, we expanded the isosurface of Duke by approximately 2 cm and constructed a Hugyens’ surface that fully surrounded the sample (Figure 12a) [11]. This basis support was discretized with 11 618 triangular elements, and a basis of incident fields was generated using RWG surface currents (see Section 5.1.5). For the coil SNR calculations, we modeled two realistic coil formers. The first former resembled a bell-shaped structure with its front region carved out (Figure 12b). The former was 24 cm long and spanned 25 cm in the x-direction and 20 cm in the y-direction. We used 9, 205 triangular elements for its discretization. The second former was an open elliptic cylinder of height *h* = 28.5 cm, semi-major axis length 14.7 cm, and semi-minor axis length 12.8 cm. We used 2 643 triangular elements for its discretization (Figure 12c). The SVD operations required for the basis generation method were performed using a threshold *ε* = 10^*−*3^ (see Section 5.1.5).

**Fig. 12:**
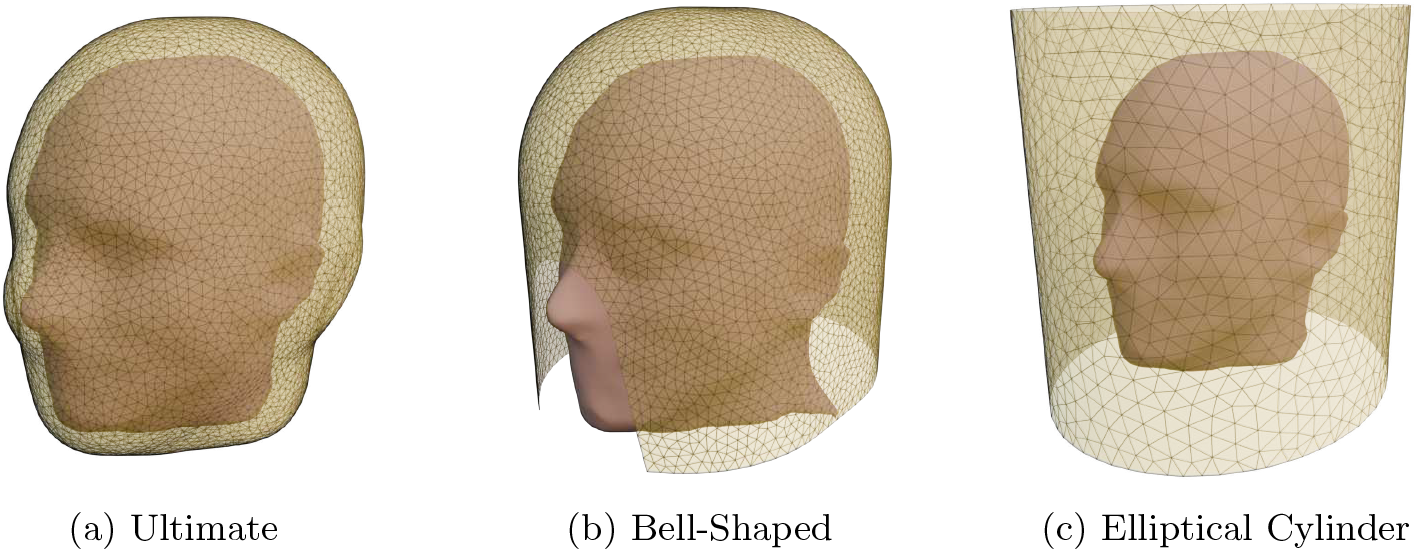
Geometry of the ultimate (a), the bell-shaped (b), the cylindrical elliptical (c) current-bearing substrate surfaces. The ultimate surface closely surrounded the realistic head model (Duke).

### 7.3 Optimization of the position of a single dipole

We optimized the position of a single dipole with length *l* = 25 cm operating at 14 T, mapped on the elliptic cylinder former (Figure 12c). The dipole was segmented with two capacitors positioned at one-quarter and three-quarters of the dipole length, which were adjusted at every step of the optimization to ensure tuning. We uniformly sampled seven voxels as the target optimization ROIs, along an elliptical trajectory on the transverse plane at *z* = 5 cm. The dipole was placed vertically to the X-Y plane. Starting from a position behind the head, we optimized the horizontal placement of the dipole on the elliptic cylinder substrate for each ROI independently. This was performed as a two-step optimization: grid search followed by a gradient-based optimization.

### 7.4 Three-coil optimization with a moving spherical ROI

We optimized a three-element array at 3 T. The loops were mapped to the bell-shaped former (Figure 12b), discretized with triangular elements of average edge length 6 mm, and segmented with seven capacitors for tuning. The values of the capacitors were adjusted to ensure tuning and ideal decoupling at every iteration of the optimization.

The optimization target was the average SNR within a spherical ROI defined on a transverse plane *z* = 5 cm with a radius *r* = 2 cm. The ROI was moved to different locations within the head (Figure 13). For location 1, we initialized the optimization with an arbitrary initial guess for the shape and position of the three coils. We then performed a four-step optimization: grid search on the coil size and position with only affine transformation, gradient-based optimization on the coil size and position, grid search on the control point position, and gradient-based optimization on the control point position (see Section 6.2). For locations 2 to 7, the initial guess was the last iteration of the second-step optimization (after affine transformation) for the previous ROI location.

**Fig. 13:**
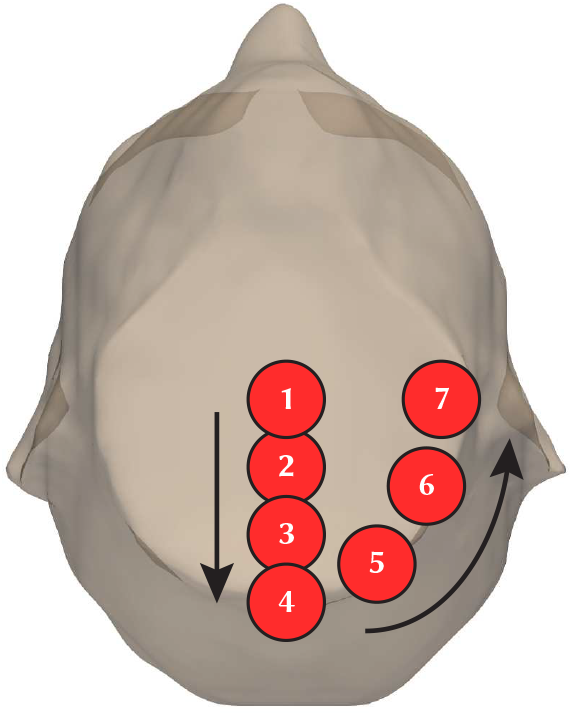
Schematic visualization of the seven locations of the spherical ROI for the three-coil optimization. All locations were centered on an axial plane at *z* = 5 cm. A manual initial guess was defined before optimizing for the ROI at location 1. For each subsequent ROI location, the coil configuration in the last iteration of the optimization for the previous location was used as initial guess.

### 7.5 Four-coil optimization with two regions-of-interest

We optimized a four-element loop array at 3 T with two separate voxels as the optimization target, one in the frontal part and the other in the rear part of the head. We also introduced a weighted formulation of the cost function in Equation (38):

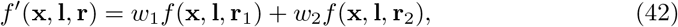

where *w*_1_ and *w*_2_ are weights that prioritize the relative importance of voxel r_1_ or r_2_ for the SNR optimization.

### 7.6 12-coil optimization

We performed three experiments aimed at optimizing a 12-coil receive array for 3 T brain imaging. We defined two ROIs: one for the cerebrum and one that combined the white matter (WM), gray matter (GM), and cerebrospinal fluid (CSF) of the Duke’s head model. We used the cost-function of equation (42). In the first experiment, we optimized the average SNR performance with equal weighting for the two ROIs; in the second experiment, we assigned a larger weight to the cerebrum; in the last experiment, we optimized the array to maximize SNR performance solely over the cerebellum. Finally, to evaluate the generalizability of the optimization process, we loaded the optimized coil configuration of the first experiment with Ella’s head model and calculated the SNR performance.

## 8 Results

### 8.1 Numerical precision of the VSIE

We used a simple loop geometry with initial node coordinates **r**_*i*_, one port, and three tuning capacitors. As the parametrization for the coordinates, we use a simple affine transformation with parameter *α*. The parametrized coordinates **r**(*α*) are defined as

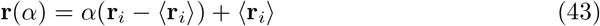

with **r**(1) = **r**_*i*_ and ⟨**r**_*i*_⟩ denoting the mean of the initial coordinates.

We estimated the gradient using the original implementation with limited precision and with the proposed implementation with enhanced precision for a number of step sizes. We estimated the gradient of the cost function about *α* = 1 using the Forward Euler approximation of the derivative, which is given by

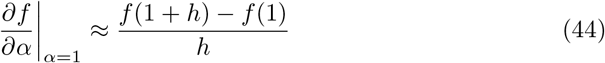

for step size *h*.

Fig. 14 shows the change in the cost function and the estimated gradient magnitude as a function of step size *h*, using the existing and proposed approaches. As can be seen in Fig. 14b, estimating the gradient of the SNR cost function fails catastrophically when the finite-difference step size is too small, even for a simple affine transformation that results in all nodes changing position. Fig. 14a shows that below *h* = 10^*−*5^, the change in the cost function becomes independent of the step size, explaining why estimating the gradient fails catastrophically.

**Fig. 14:**
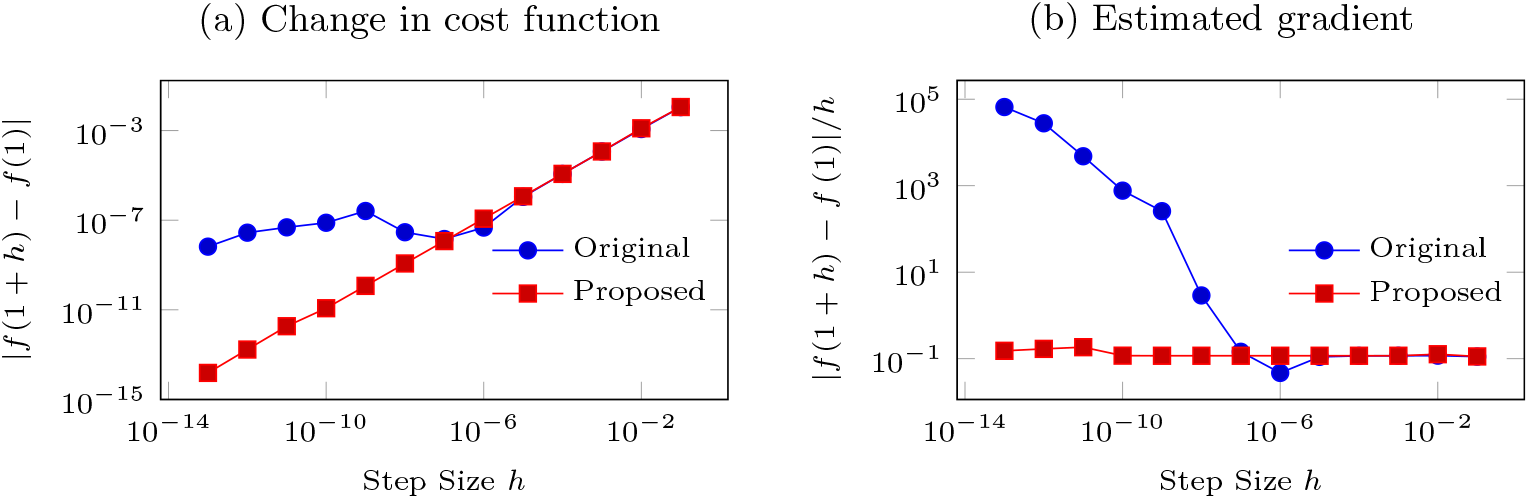
Change in cost function and estimated gradient of cost function about *α* = 1 as a function of Forward Euler finite difference step size using original implementation and proposed implementation.

### 8.2 Coil design optimization experiments

Without the MRGF, the iterative optimization process would have been intractable. For example, the MRGF approach resulted in a solution time, including tuning and decoupling, of 15 s for each three-coil design, as opposed to a few minute when assembling the full system in Equation (16). The full solution would take several minutes for the 12-coil array, whereas it required 32 s with MRGF. The optimization of the dipole position required seven minutes for each ROI location. The length of the full coil optimization ranged between 5 and 21 hours for the seven three-coil optimization experiments (Section 7.4), and between a week and two weeks for the four 12-coil optimization experiments (Section 7.6). Note that the exact duration of the individual iterations is not reported since it varied during the optimization based on the convergence of each step of the grid search and gradient-based optimization.

Figure 15 shows that the optimization algorithm adapts the position of the dipole as the position of the target ROI moves around the head. Since in this experiment we did not optimize the shape of the coil but only its azimuthal position, the relative SNR performance of the dipole was lower for ROI 6 and ROI 7 because they are closer to the surface of the head, where the UISNR grows exponentially.

**Fig. 15:**
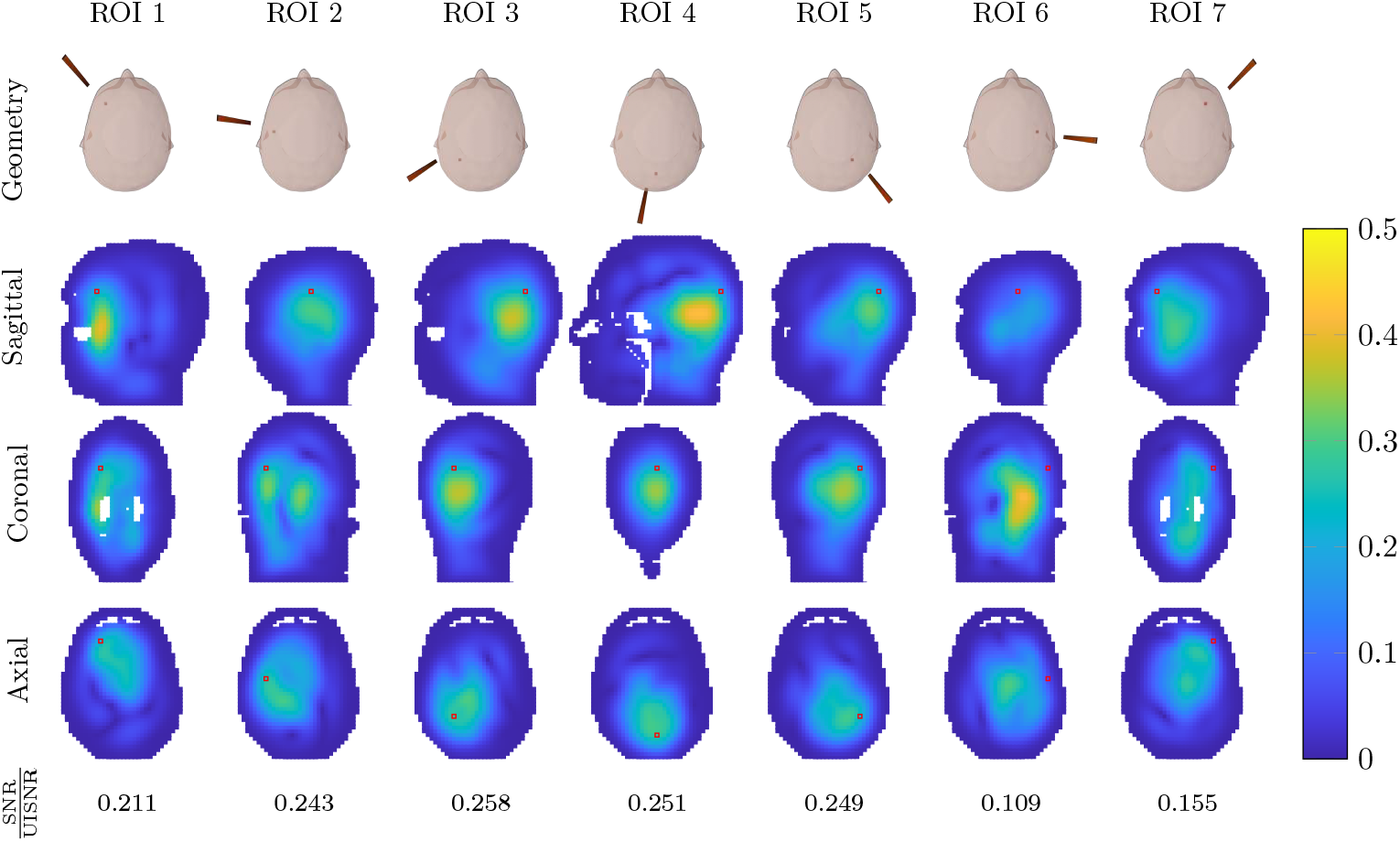
Optimization of the absolute SNR performance of a dipole at 600 MHz (14 T), as the target ROI is moved around the head. The dipole was constrained to move azimuthally on a cylindrical former surrounding the head. From top to bottom row: position of the dipole resulting from the optimization, with the red voxel indicating the target ROI; performance maps showing the percentage of the UISNR achieved by the dipole along sagittal, coronal, and axial sections cutting through the target voxel (in red). The absolute performance for each configuration is reported at the bottom.

Figures 17–30 show the results of the three-coil optimization for the seven positions of the spherical ROI.

For each experiment, there are two figures. The first figure shows the evolution of the cost function during the four steps of the optimization: grid search on the coil size and position with only affine transformation (Grid Search 1), gradient-based optimization on the coil size and position (Gradient-based optimization 1), grid search on the control point position (Grid Search 2), and gradient-based optimization on the control point position (Gradient-based optimization 2). The second figure shows the evolution of the coil design for different iterations, together with the associated performance maps for three orthogonal section of the head cutting through the center of the spherical ROI. In these and subsequent figures, the coils were constrained to be on a bell-shaped former surrounding the head. In each figure, from top to bottom, the rows show: geometry and ROI (in red); performance maps showing the percentage of the UISNR achieved by each configuration along sagittal, coronal, and axial sections cutting through the center of the ROI (contour shown in red). The mean and maximum performance within the ROI is reported at the bottom.

For all cases, the average ratio of SNR over the UISNR within the ROI increased monotonically with the number of iterations. The performance was lower when the ROI was closer to the surface of the head, where it is more difficult to approach the UISNR [10]. Since the three coils were constrained to the bell-shaped former, the performance difference among ROI positions was also affected by the distance between the former and the surface of the head. Table 2 summarizes the results of the three-coil experiments. Except for ROI 1, for which the optimization was initialized to a random geometry, in all other cases the initial guess was the coil geometry optimized for the previous ROI position. The values in Table 2 show that the cost function decreased in each case. The optimization algorithm converged to a different coil geometry as the ROI moved to a different position. Figure 16 shows that the three-coil geometry optimized for ROI 4 is consistent with the corresponding ideal current patterns associated with the UISNR.

**Table 2:** Summary of three-coil optimization results. For each position of the spherical ROI (Figure 13), we report: the cost function of the initial guess, the cost function at the end of the optimization, the average and maximum performance within the ROI after the optimization.

| ID | Cost Function<br>(first iteration)<br>% | Cost Function<br>(final iteration)<br>% | Average<br>SNR/UISNR<br>% | Maximum<br>SNR/UISNR<br>% |
| --- | --- | --- | --- | --- |
| ROI 1 | 64.4 | 26.4 | 74.7 | 83.9 |
| ROI 2 | 35.7 | 23.2 | 77.9 | 88.0 |
| ROI 3 | 38.5 | 29.8 | 73.2 | 86.3 |
| ROI 4 | 60.9 | 54.8 | 54.5 | 79.7 |
| ROI 5 | 63.8 | 57.7 | 52.8 | 79.0 |
| ROI 6 | 72.5 | 65.9 | 46.7 | 75.2 |
| ROI 7 | 84.2 | 80.2 | 34.1 | 66.2 |

**Fig. 16:**
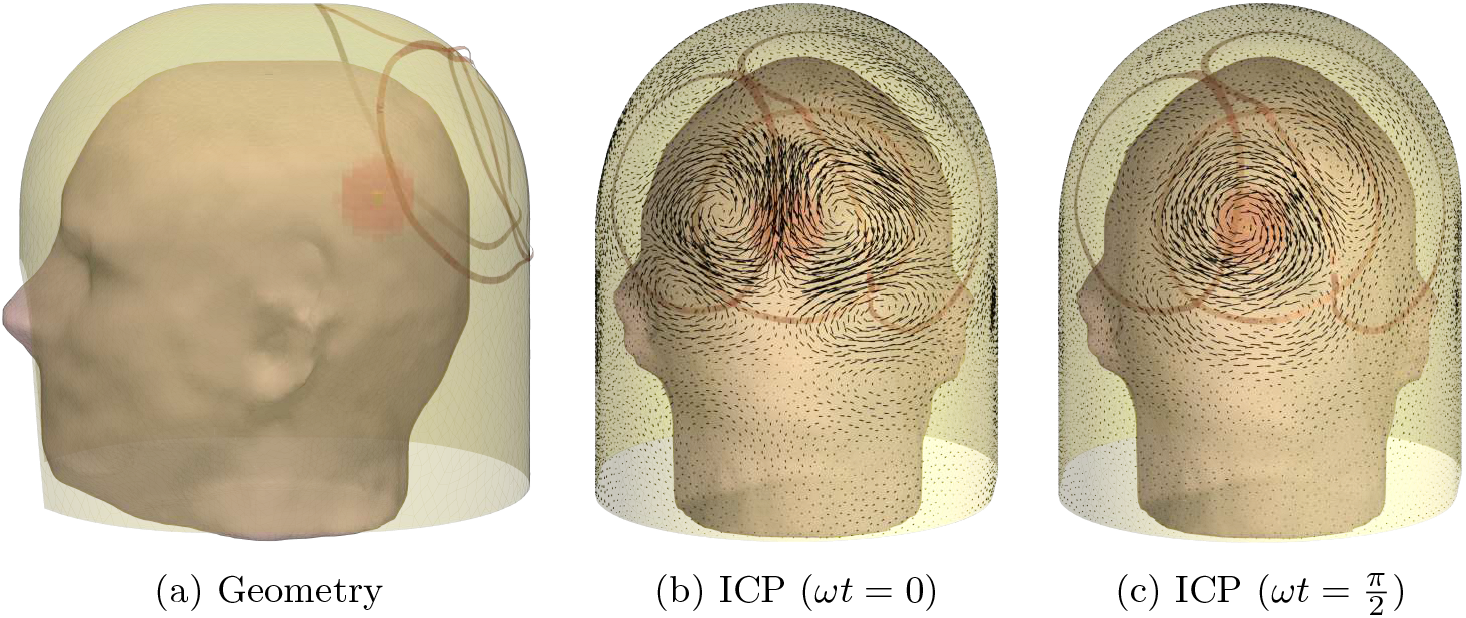
Realistic coil former surrounding the head model along with optimal coil configuration for spherical ROI 4 (a). The former also served as a current-bearing surface for the estimation of the ideal current patterns (ICP). Temporal snapshots of the ICP yielding optimal signal-to-noise ratio at the center of the spherical ROI 4 for two time-points are shown in (b) and (c).

**Fig. 17:**
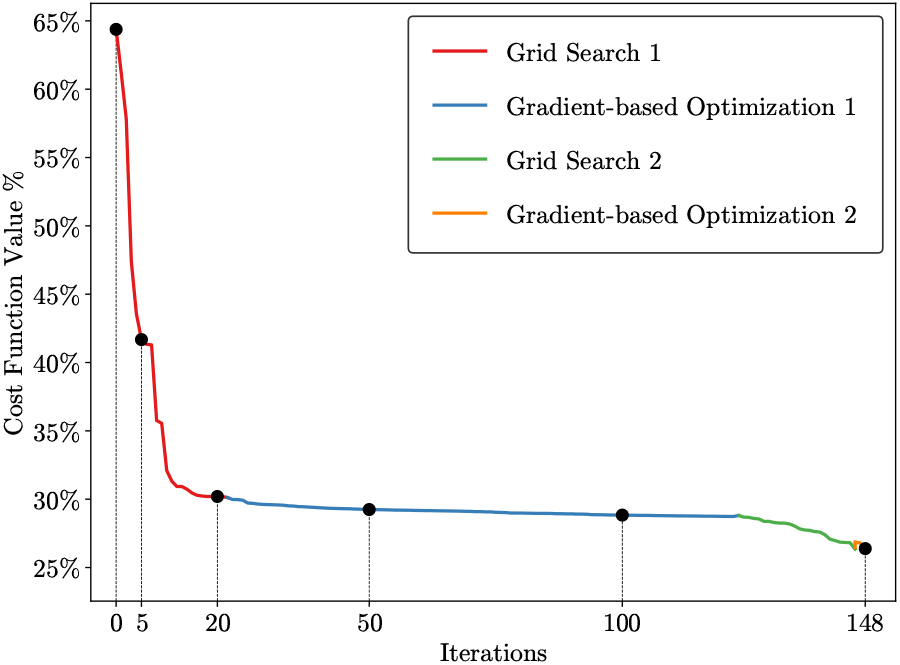
The cost function evolution for three-coil optimization with the spherical ROI in region 1.

**Fig. 18:**
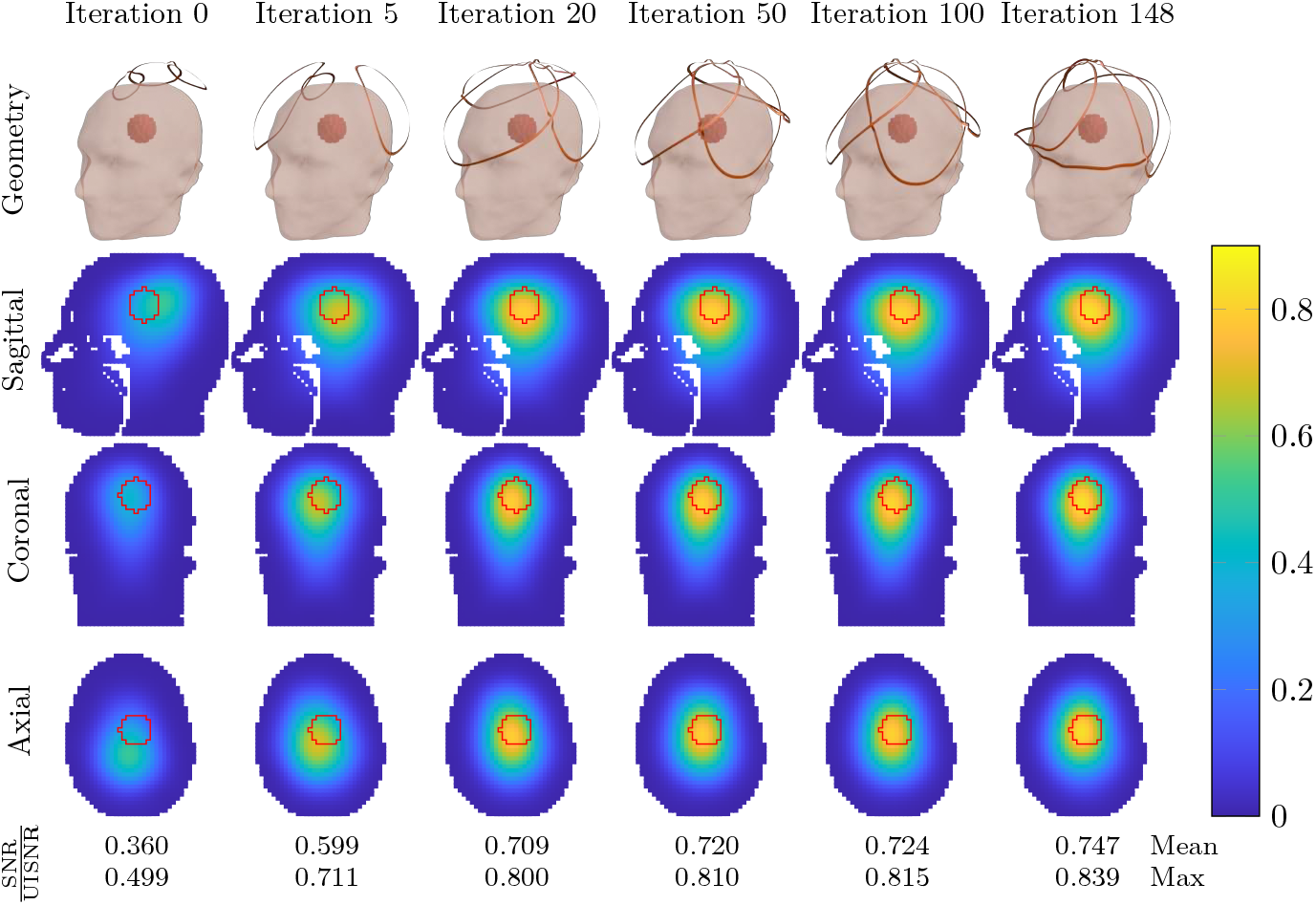
Evolution of the three-coil geometry during the iterative optimization of the average SNR performance within the spherical ROI in region 1.

**Fig. 19:**
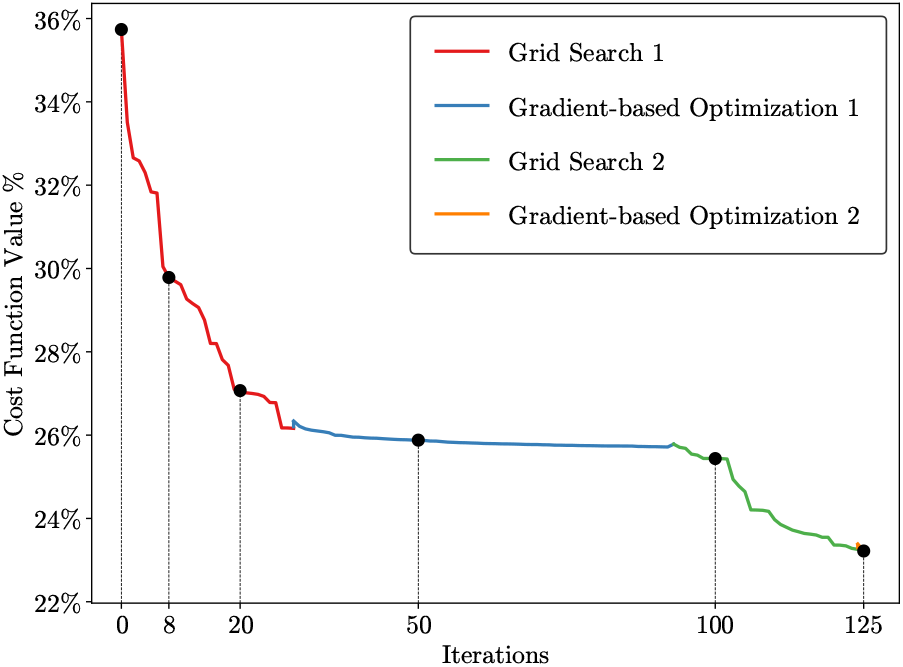
The cost function evolution for three-coil optimization with the spherical ROI in region 2.

**Fig. 20:**
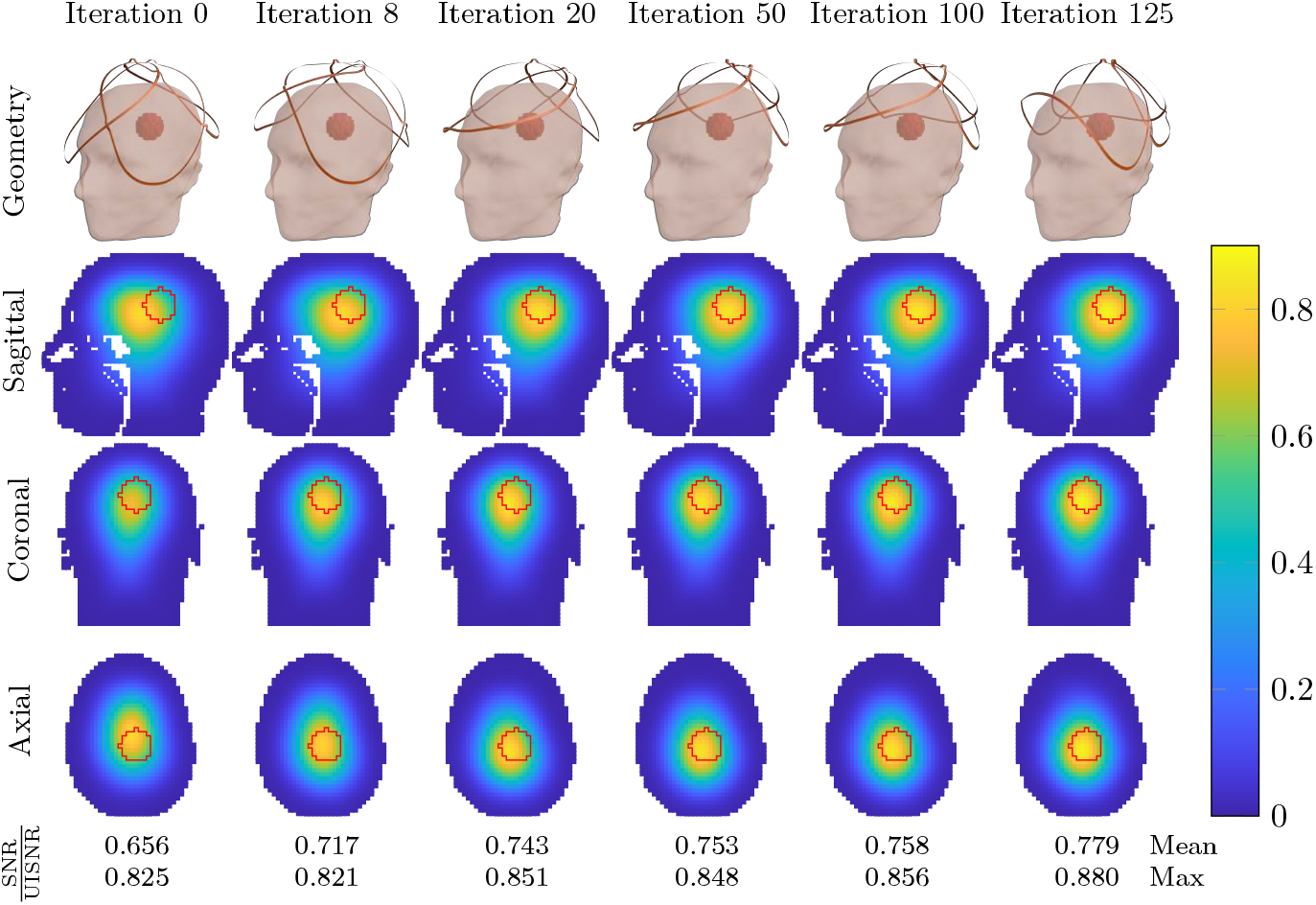
Evolution of the three-coil geometry during the iterative optimization of the average SNR performance within the spherical ROI in region 2.

**Fig. 21:**
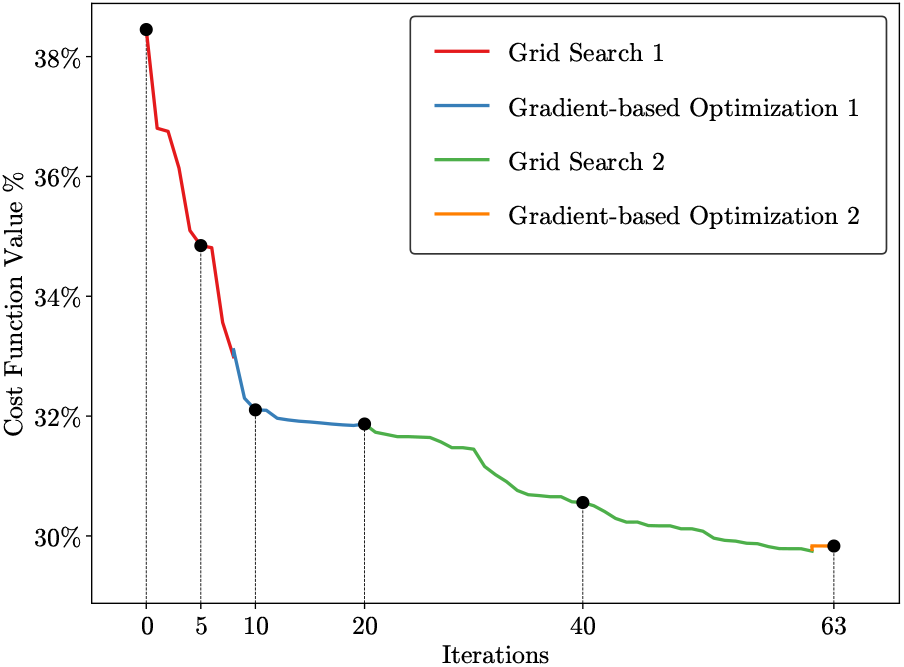
The cost function evolution for three-coil optimization with the spherical ROI in region 3.

**Fig. 22:**
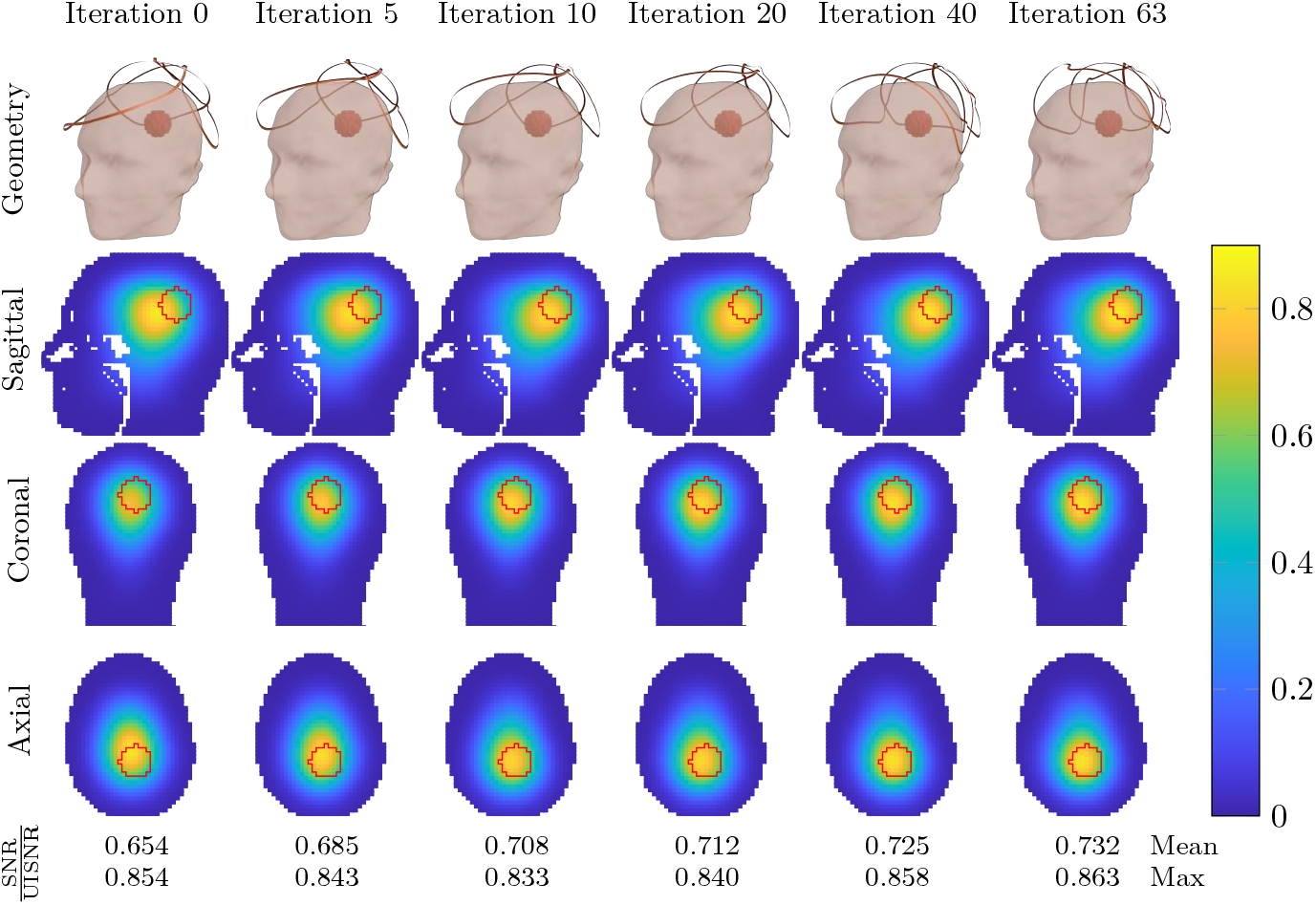
Evolution of the three-coil geometry during the iterative optimization of the average SNR performance within the spherical ROI in region 3.

**Fig. 23:**
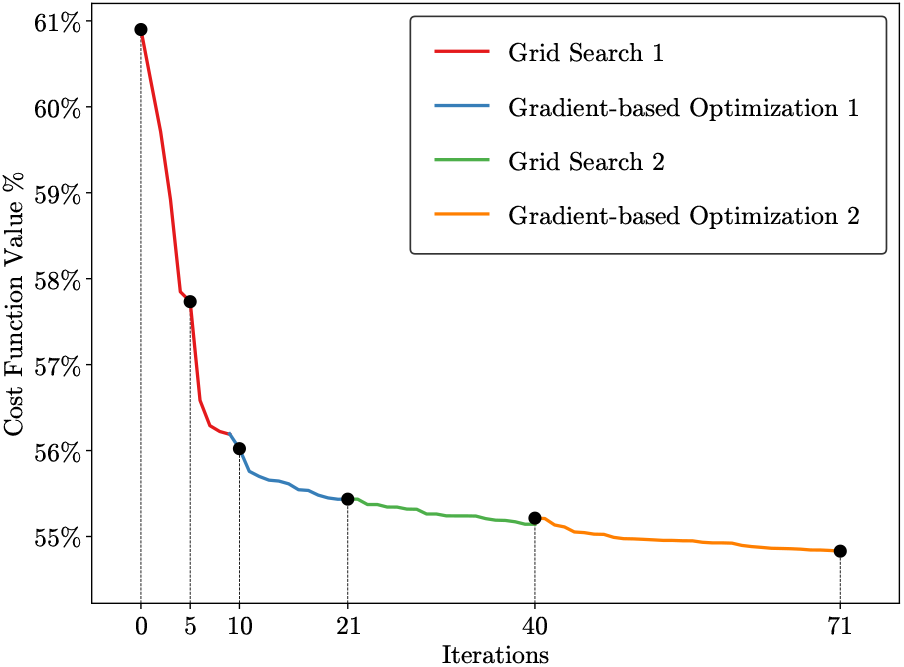
The cost function evolution for three-coil optimization with the spherical ROI in region 4.

**Fig. 24:**
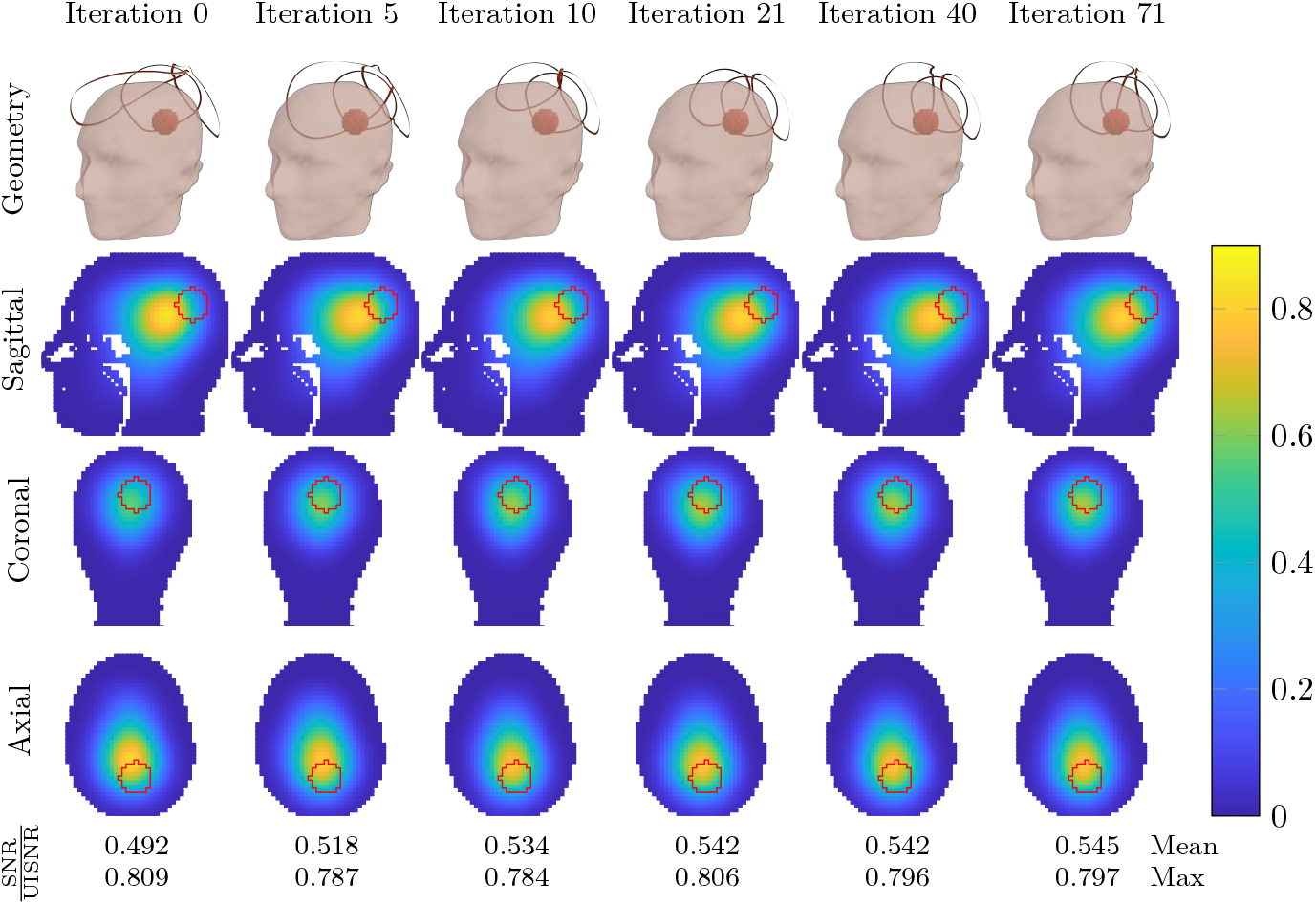
Evolution of the three-coil geometry during the iterative optimization of the average SNR performance within the spherical ROI in region 4.

**Fig. 25:**
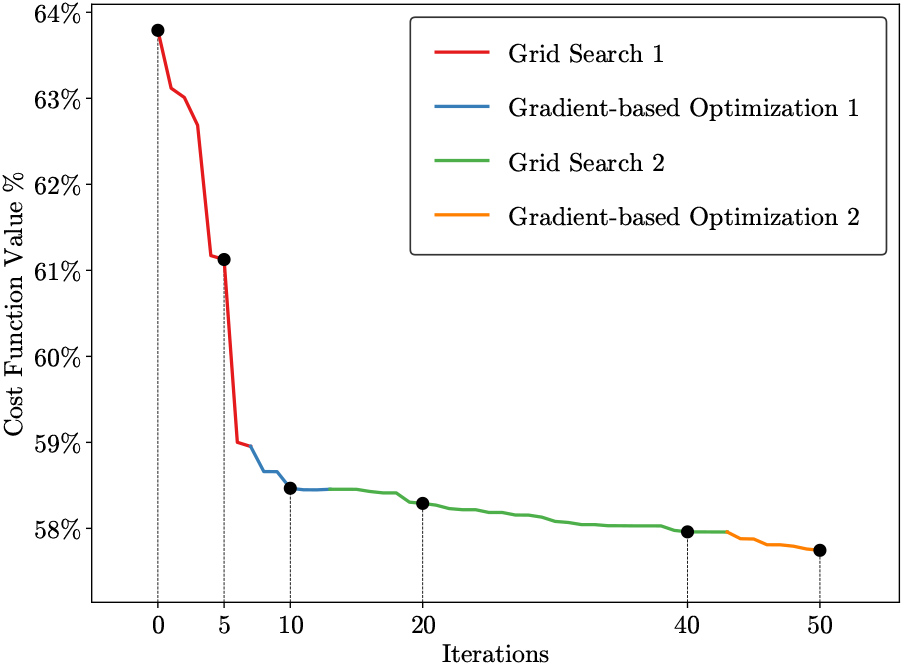
The cost function evolution for three-coil optimization with the spherical ROI in region 5.

**Fig. 26:**
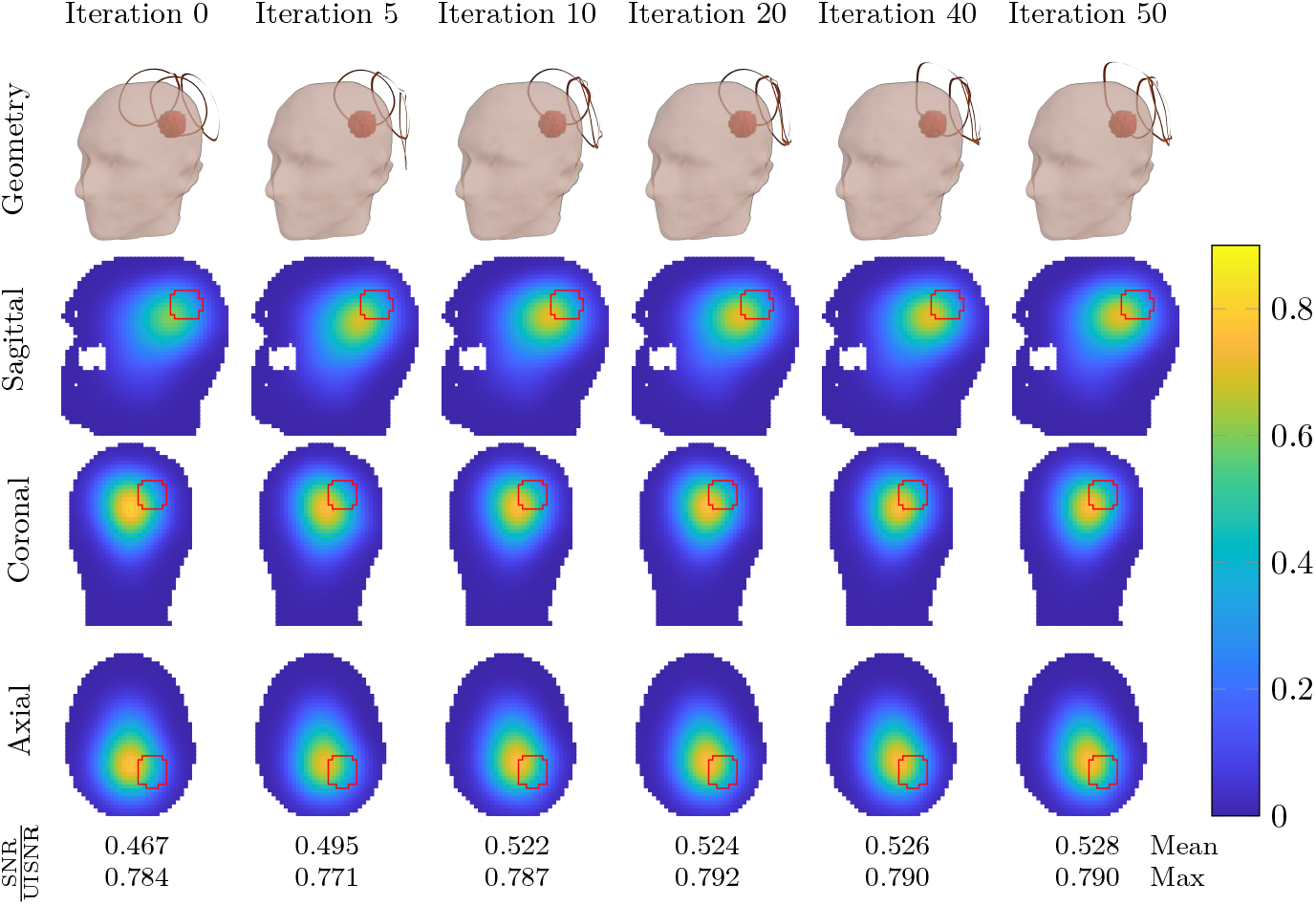
Evolution of the three-coil geometry during the iterative optimization of the average SNR performance within the spherical ROI in region 5.

**Fig. 27:**
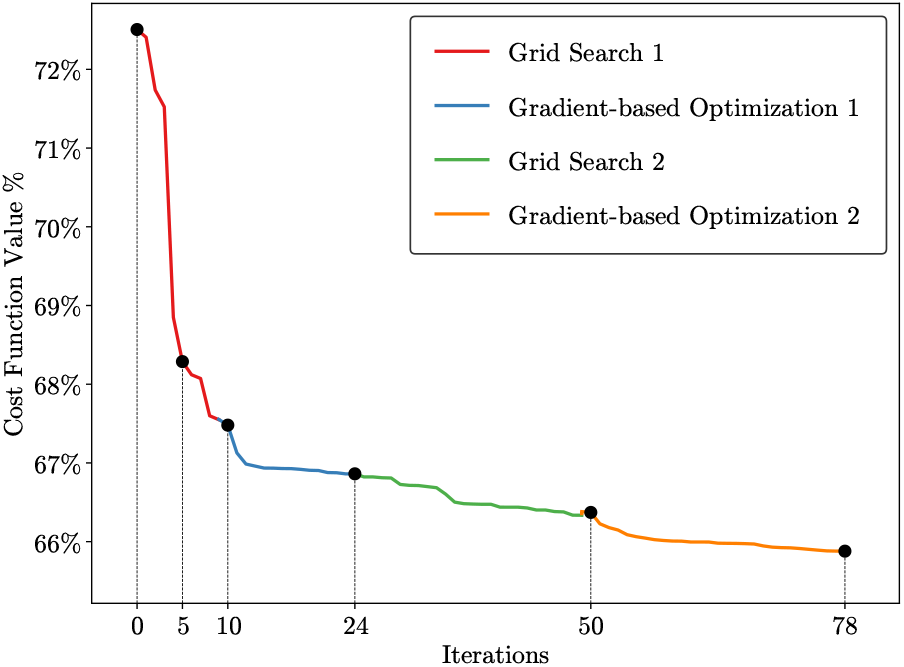
The cost function evolution for three-coil optimization with the spherical ROI in region 6.

**Fig. 28:**
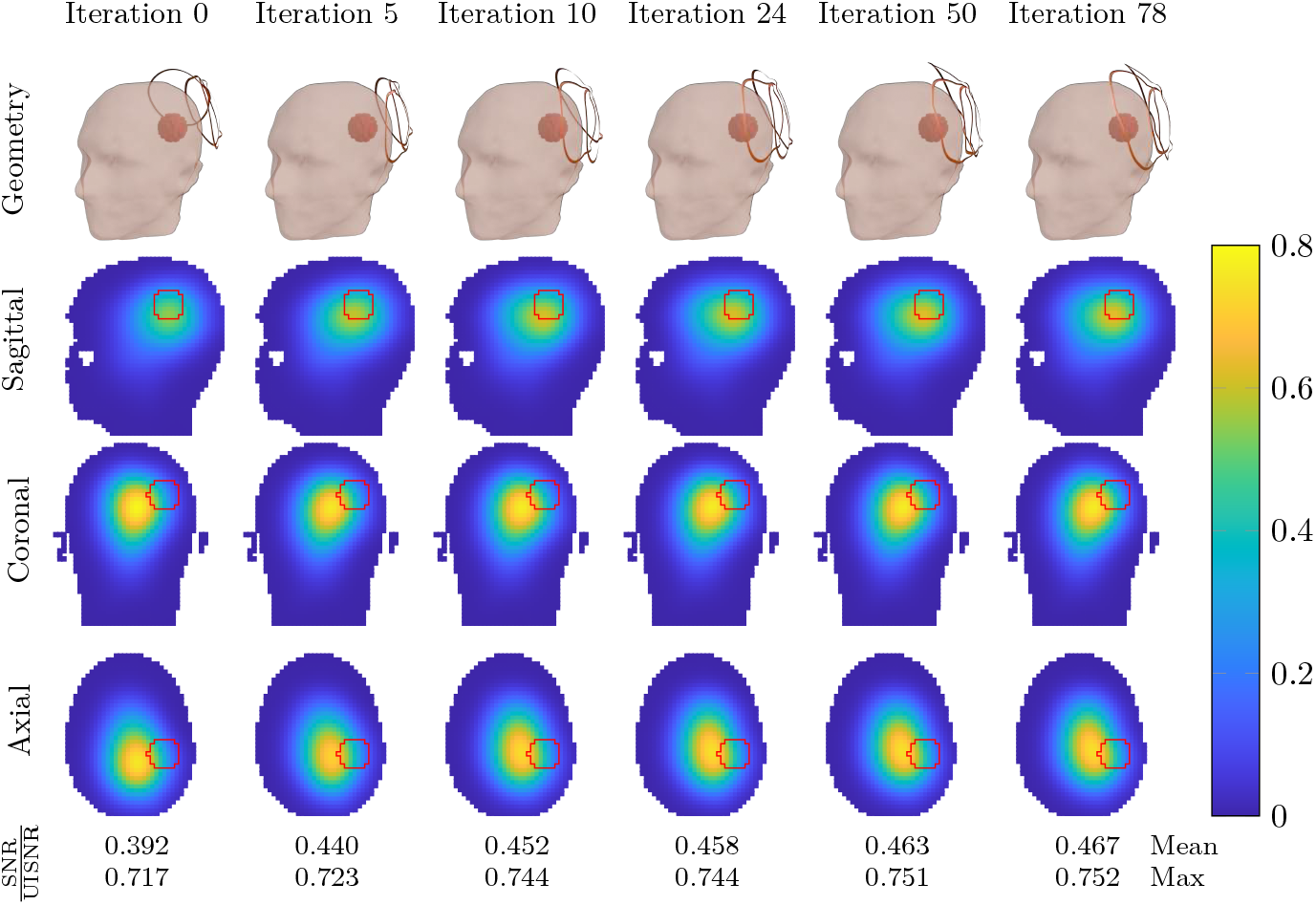
Evolution of the three-coil geometry during the iterative optimization of the average SNR performance within the spherical ROI in region 6.

**Fig. 29:**
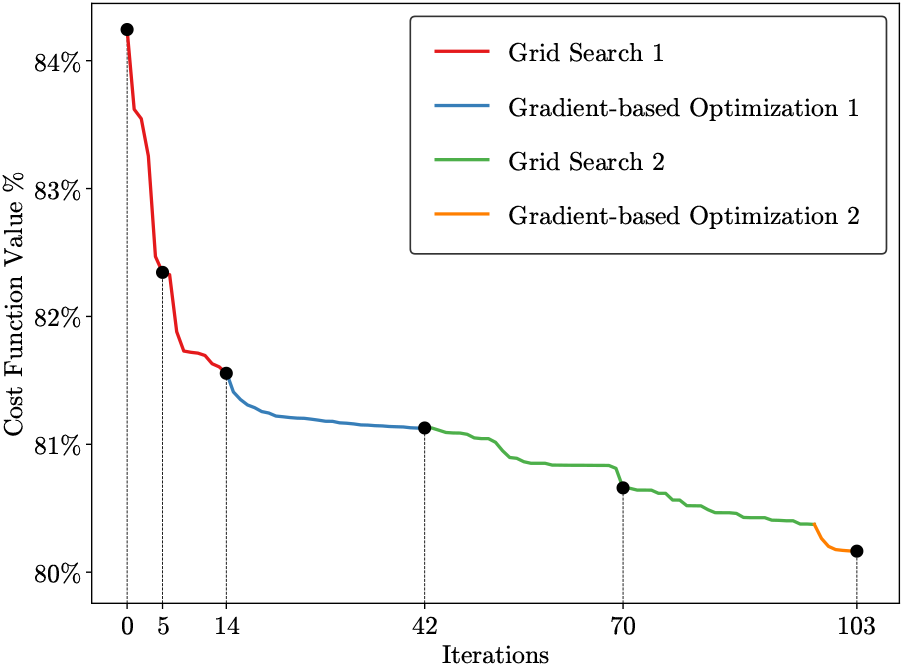
The cost function evolution for three-coil optimization with the spherical ROI in region 7.

**Fig. 30:**
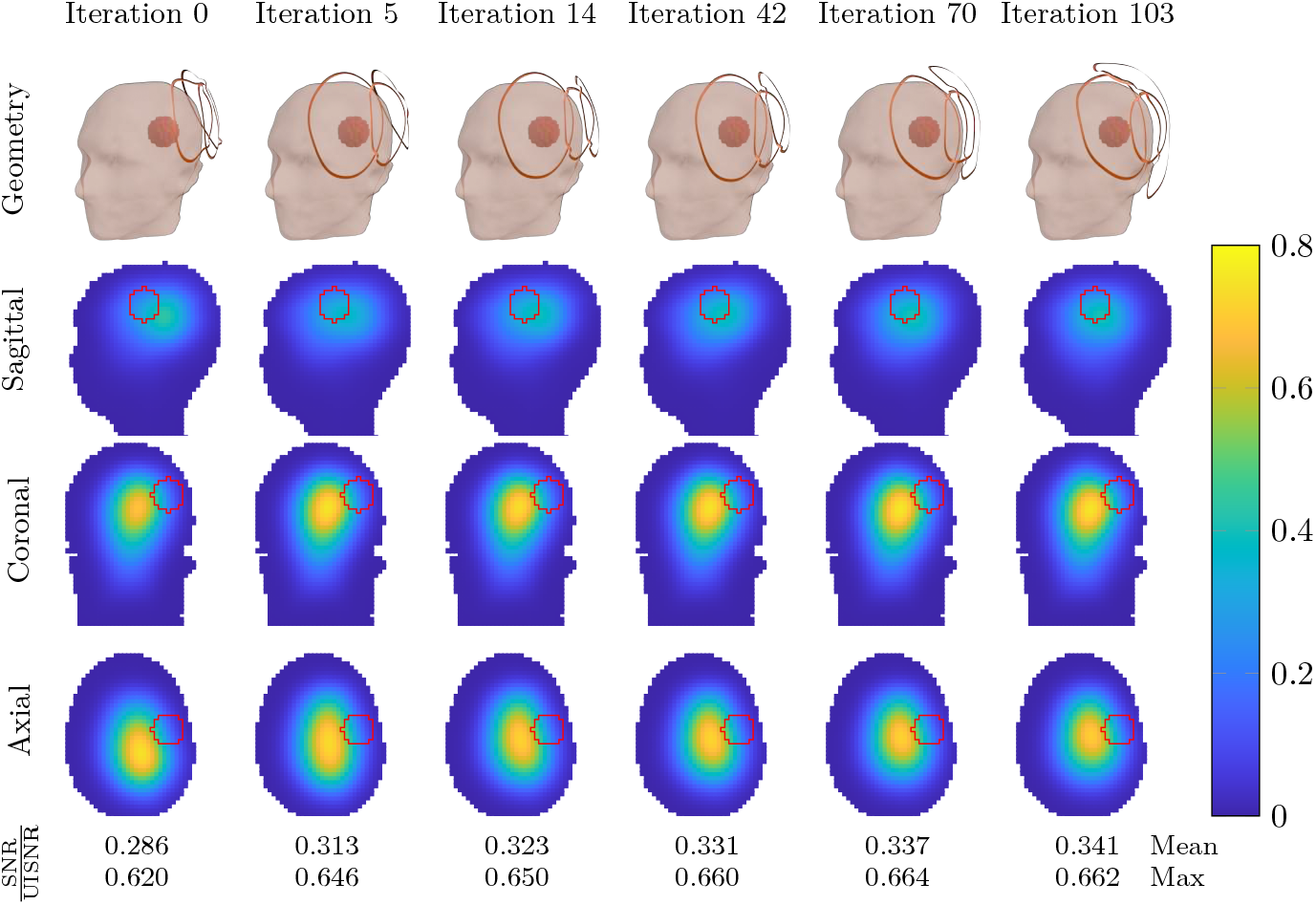
Evolution of the three-coil geometry during the iterative optimization of the average SNR performance within the spherical ROI in region 7.

Using Equation (38), the four-coil optimization starting from an arbitrary initial guess converged to a geometry in which three coils were clustered near the target voxel at the back of the head and one coil was located near the target voxel in the front (Figures 31 and 32). When the target voxels were separately considered, using Equation (42) with *w*_1_ = *w*_2_ = 0.5, the optimized configuration yielded two coils in the front of the head and two coils at the back (Figures 33 and 34). The optimization with the weighted cost function achieved higher SNR performance than the one that considered both voxels as a single target ROI.

**Fig. 31:**
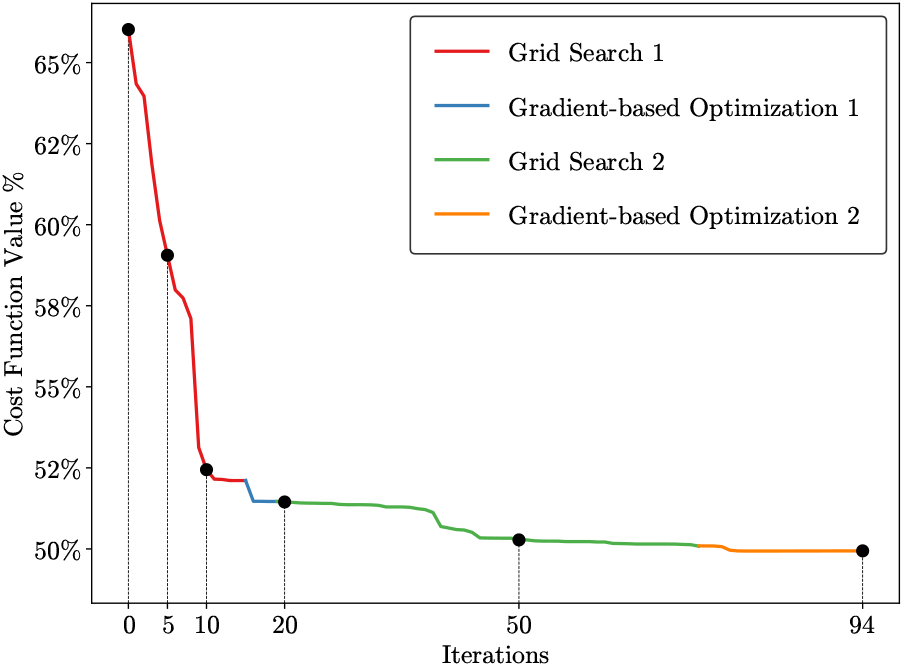
The cost function evolution for four-coil optimization with a two-voxel ROI.

**Fig. 32:**
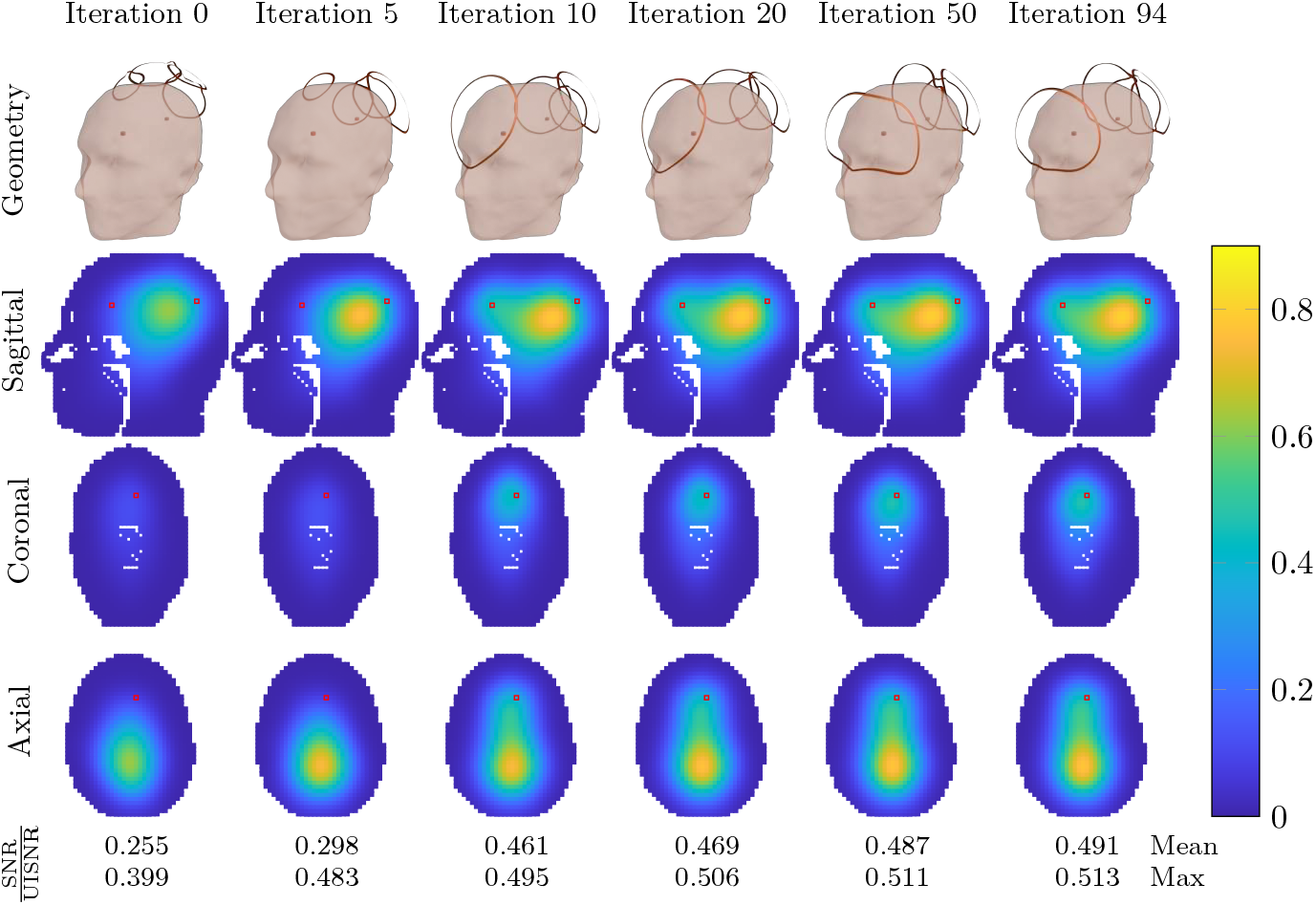
Evolution of the four-coil geometry during the iterative optimization of the average SNR performance within a two-voxel ROI.

**Fig. 33:**
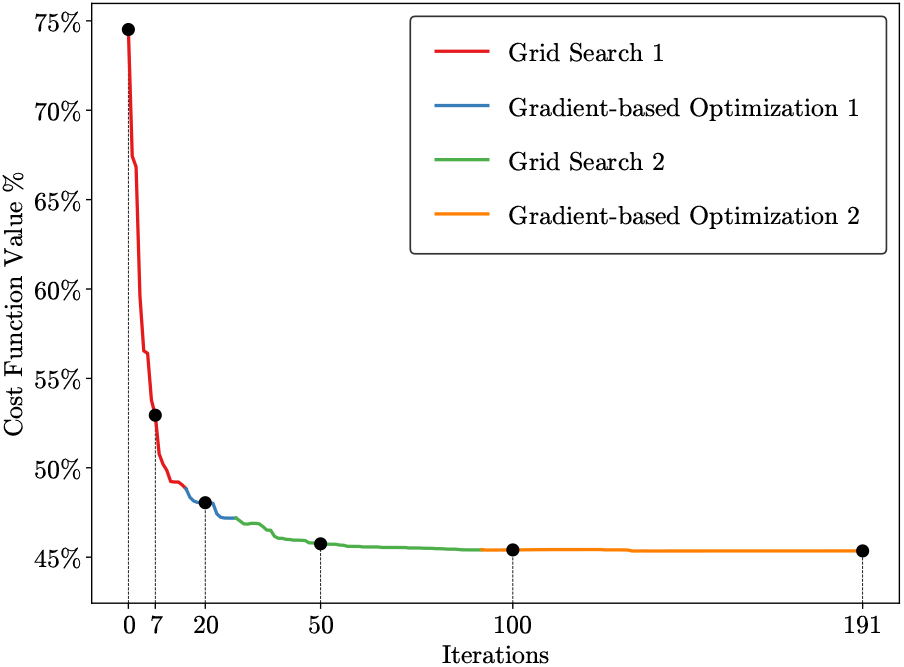
The cost function evolution for four-coil optimization with two target voxels, equally weighted and considered as separate ROIs.

**Fig. 34:**
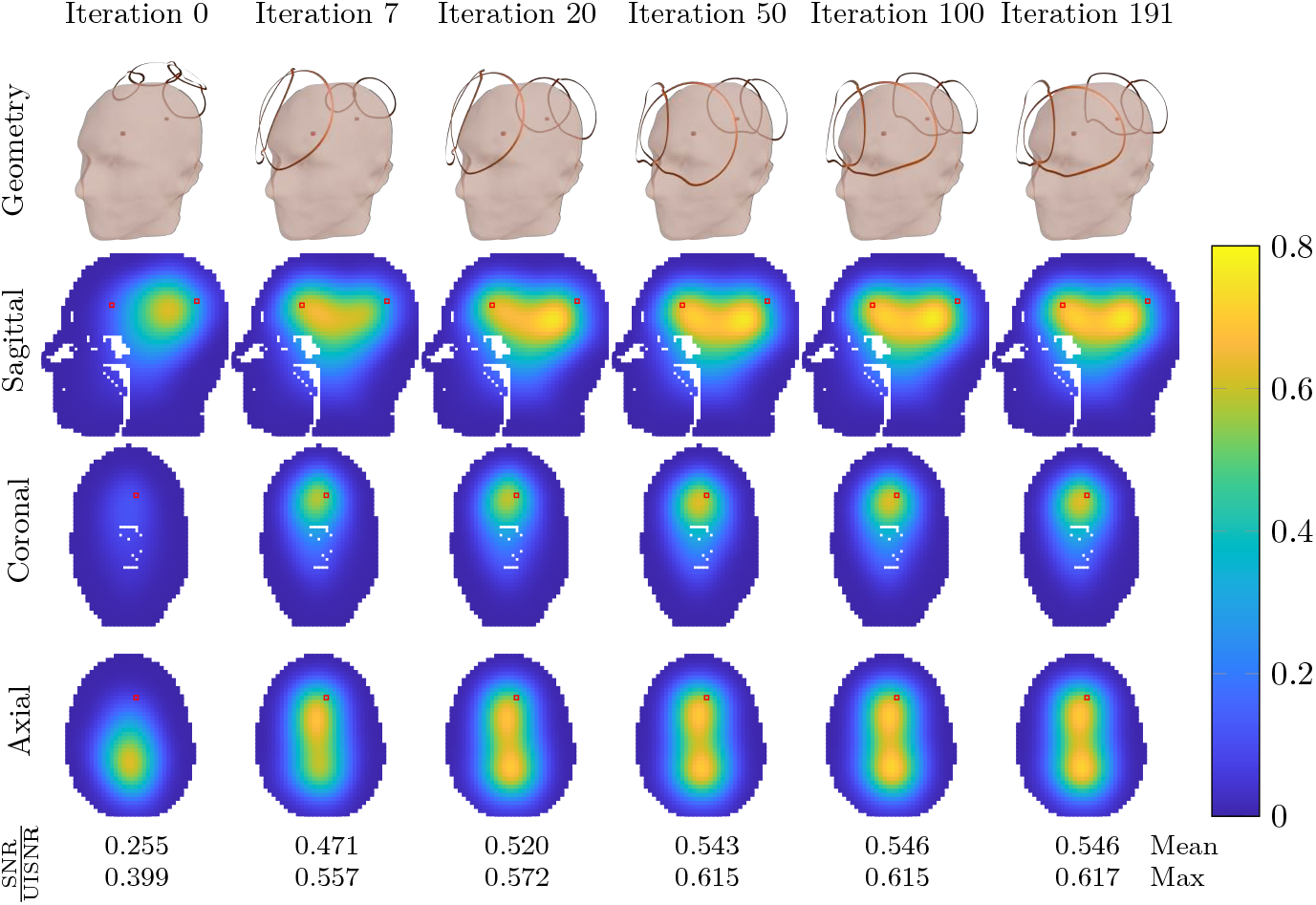
Evolution of the four-coil geometry during the iterative optimization of the average SNR performance within two target voxels, equally weighted and considered as separate ROIs.

For all 3 T 12-coil array experiments, we used an arbitrary initial guess with loop elements surrounding the head. Figures 35 and 36 show the results using equal weights *w*_1_ = *w*_2_ = 0.5 for the two ROIs. The optimized coil design achieved 7% higher performance at the center of the head and improved the averaged SNR performance over the entire brain (both ROIs) by 9%. The average performance in the cerebrum and WM-GM-CSF was 57.8% and 47.4%, respectively. Figures 37 and 38 show the optimization results when weights *w*_1_ = 0.6 and *w*_2_ = 0.4 are used for the cerebellum and WM-GM-CSF, respectively. The optimized SNR performance in the center of the head was 6% higher than the initial guess, and the averaged SNR performance over 35 the entire brain improved by 8%. The average performance in the cerebrum and WM-GM-CSF was 57.5% and 46.5%, respectively. Figures 39 and 40 show the results when optimizing for the cerebellum alone (*w*_1_ = 1, *w*_2_ = 0). The region with the highest performance shifted to the center of the cerebellum, where the performance improved on average by 14%. The average SNR performance in the cerebrum and WM-GM-CSF was 59.2% and 27.2%, respectively. Note that this example was included only for completeness, because it would neither be practical nor useful to construct a 12-coil array to image exclusively the cerebellum. Figures 41 and 42 show the results when the cerebellum ROI was excluded from the optimization target (*w*_1_ = 0, *w*_2_ = 1). In this case, the optimized coil design achieved 8.5% higher performance at the center of the head and improved the averaged SNR performance over the entire brain by 10%. The average SNR performance in the cerebrum and WM-GM-CSF was 42.2% and 48.9%, respectively.

**Fig. 35:**
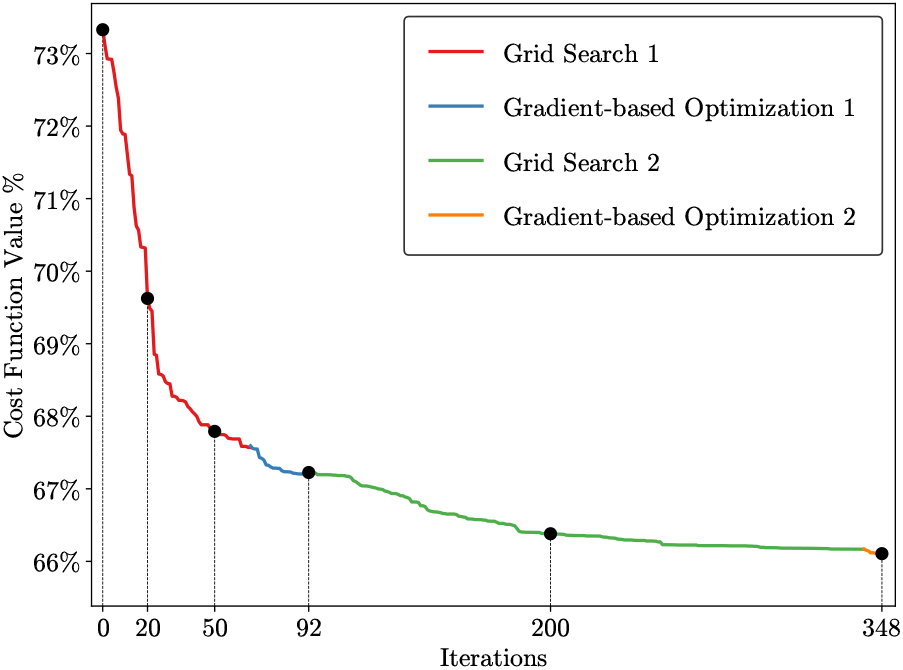
The cost function evolution for 12-coil optimization with the cerebellum and cerebrum ROIs, equally weighted (*w*_1_ = *w*_2_ = 0.5).

**Fig. 36:**
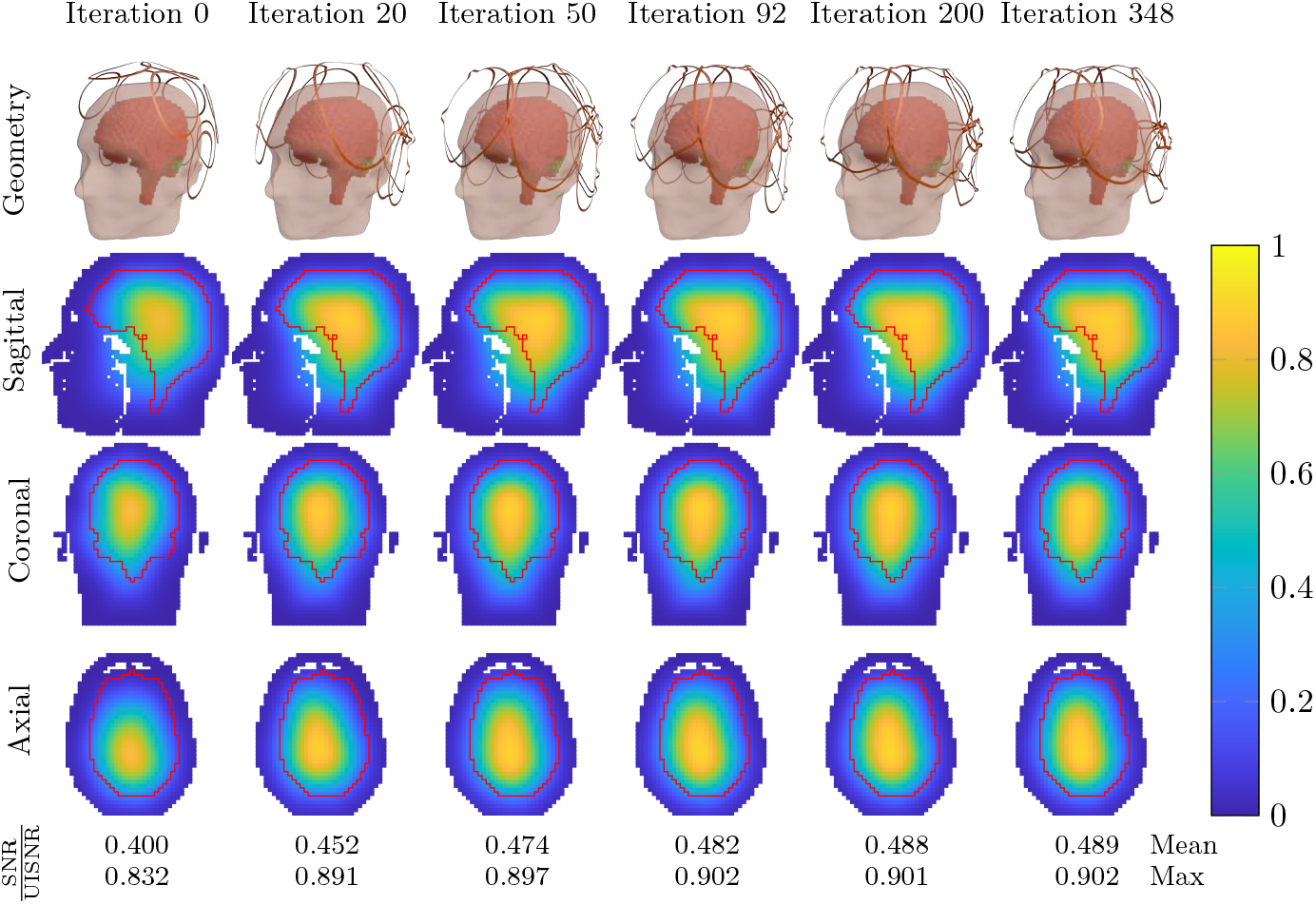
Evolution of the 12-coil geometry during the iterative optimization of the average SNR performance within the cerebellum and cerebrum ROIs, equally weighted (*w*_1_ = *w*_2_ = 0.5).

**Fig. 37:**
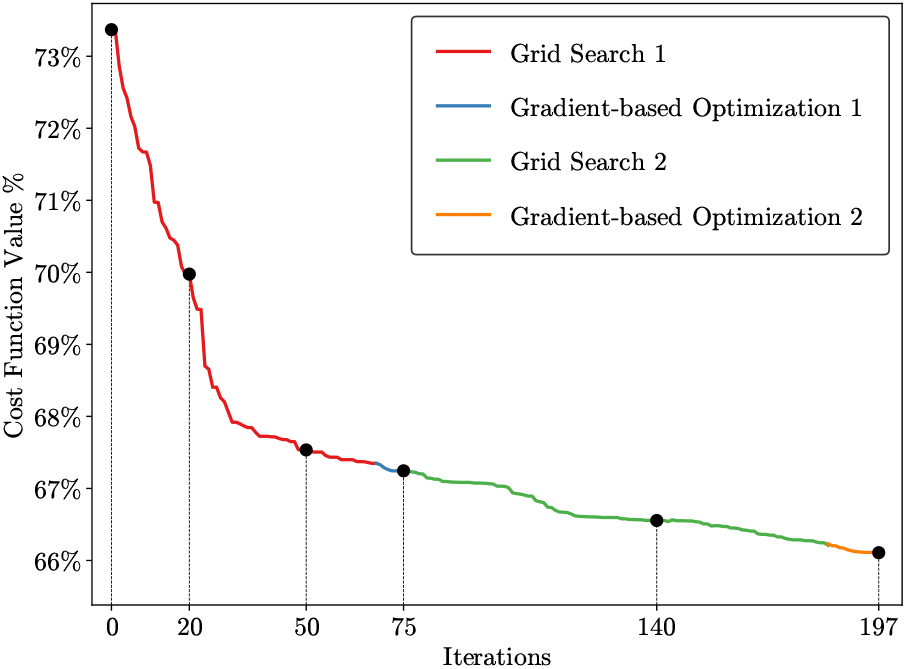
The cost function evolution for 12-coil optimization with the cerebellum and cerebrum ROIs, with weights *w*_1_ = 0.6, *w*_2_ = 0.4.

**Fig. 38:**
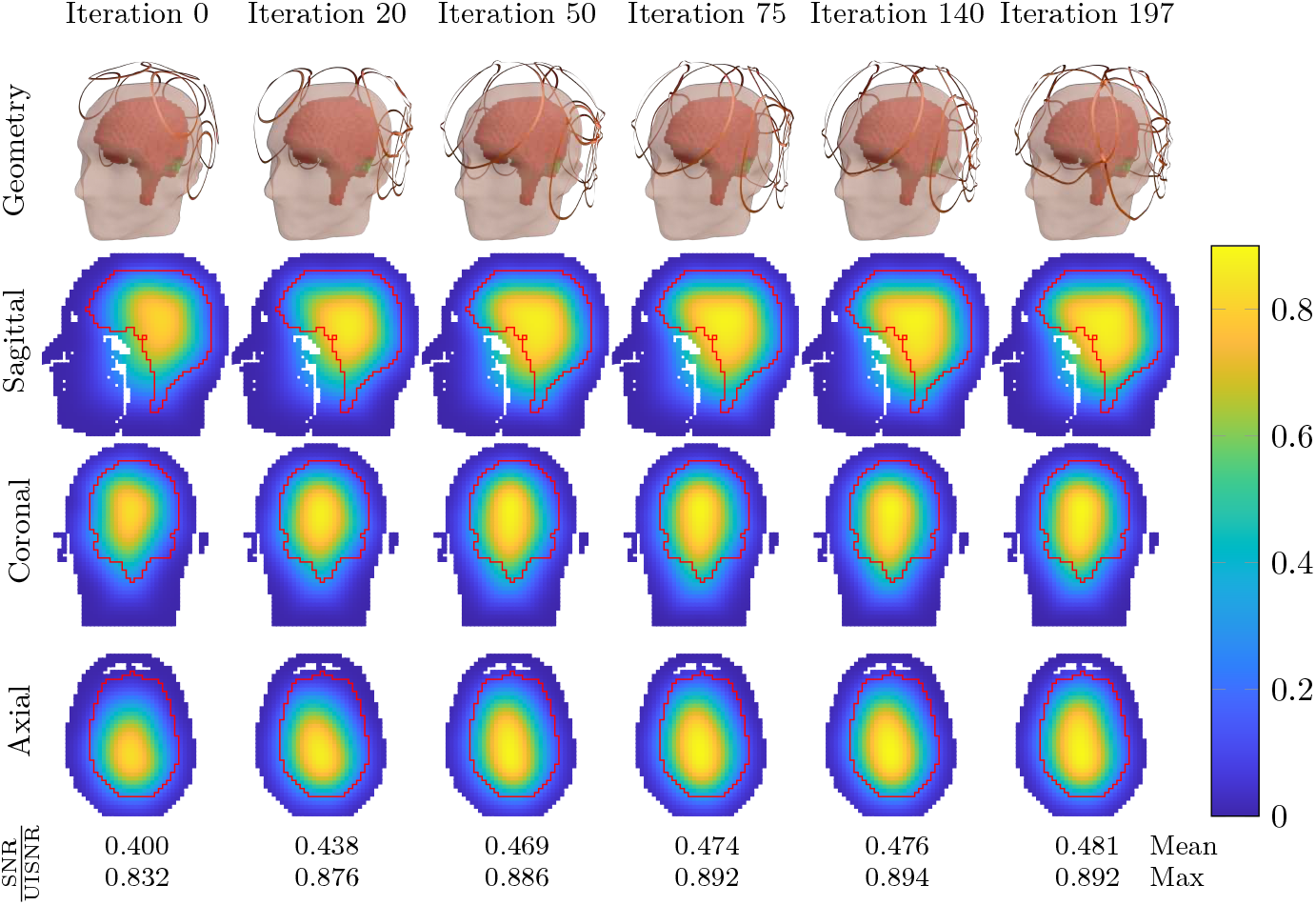
Evolution of the 12-coil geometry during the iterative optimization of the average SNR performance within the cerebellum and cerebrum ROIs, with weights *w*_1_ = 0.6, *w*_2_ = 0.4.

**Fig. 39:**
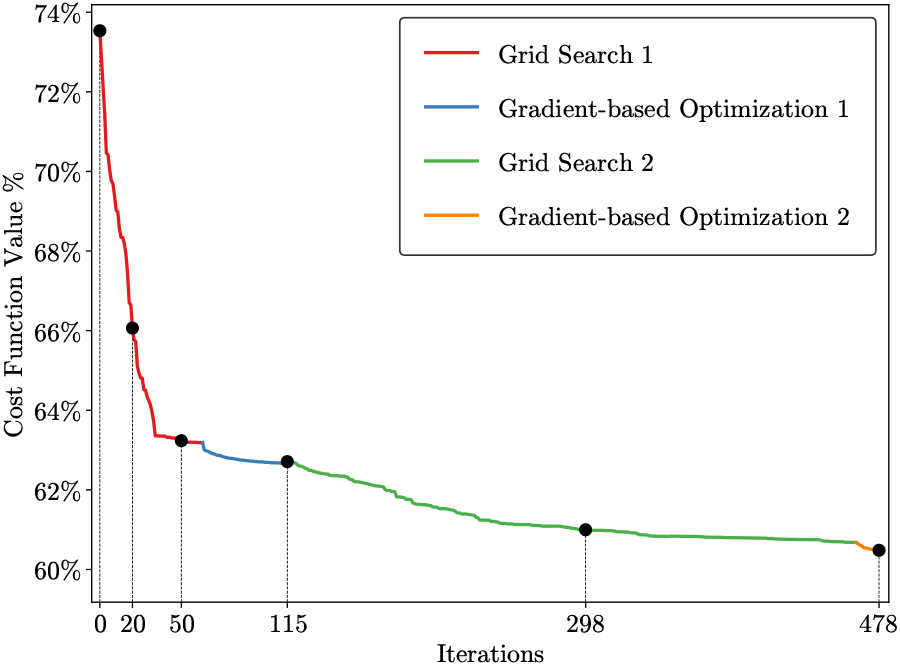
The cost function evolution for 12-coil optimization with the cerebellum ROI.

**Fig. 40:**
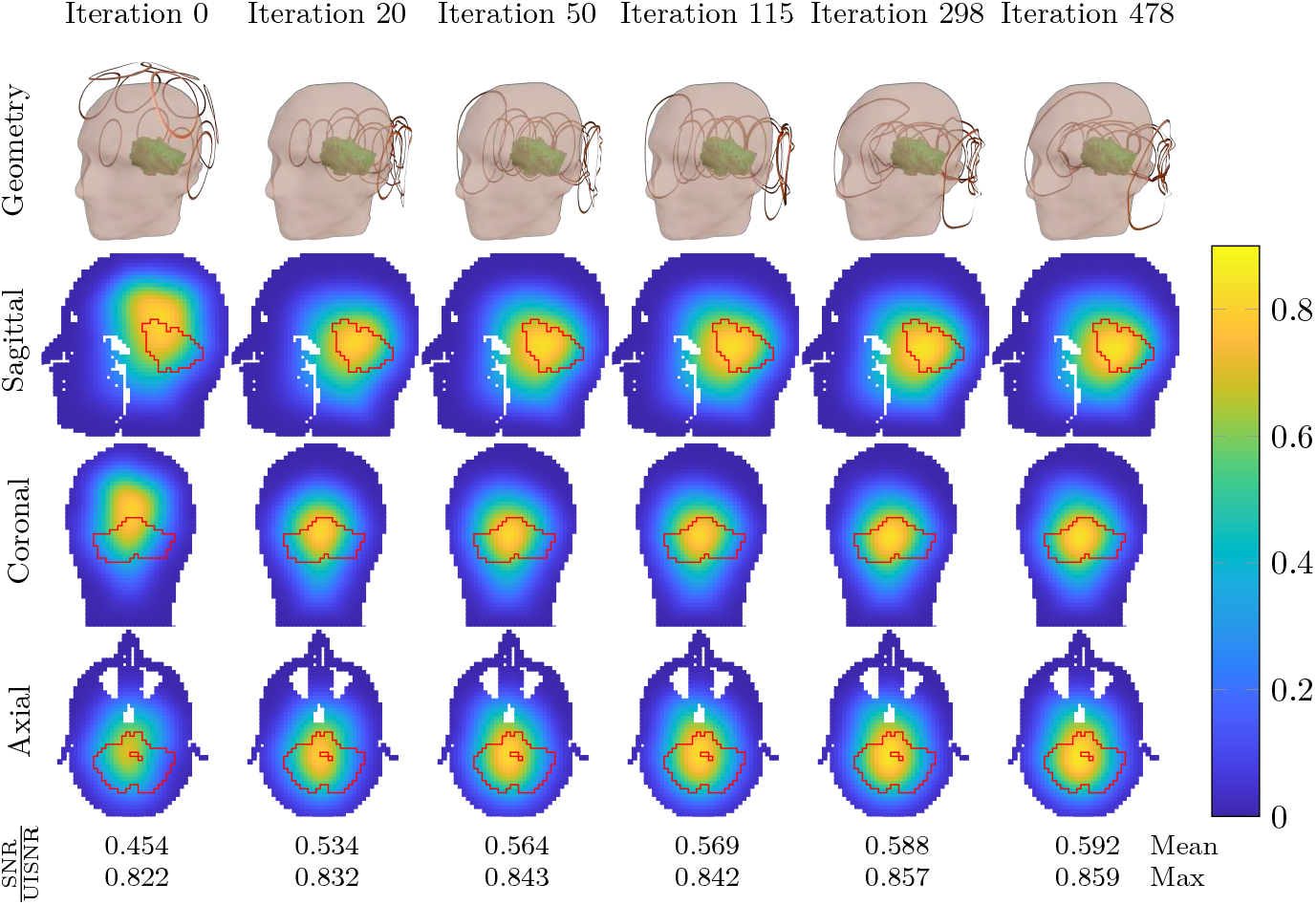
Evolution of the 12-coil geometry during the iterative optimization of the average SNR performance within the cerebellum ROI.

**Fig. 41:**
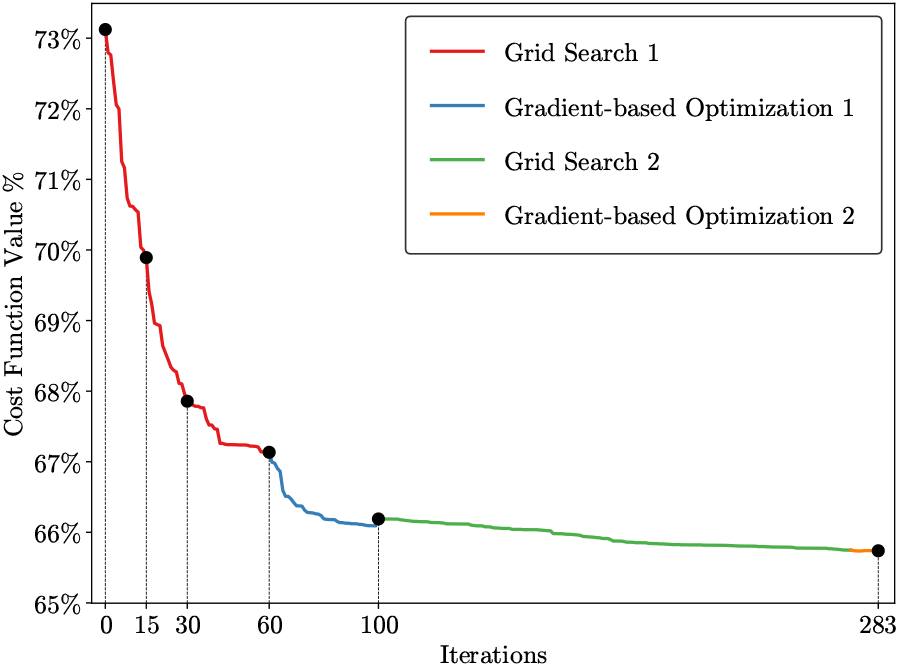
The cost function evolution for 12-coil optimization with the cerebrum ROI.

**Fig. 42:**
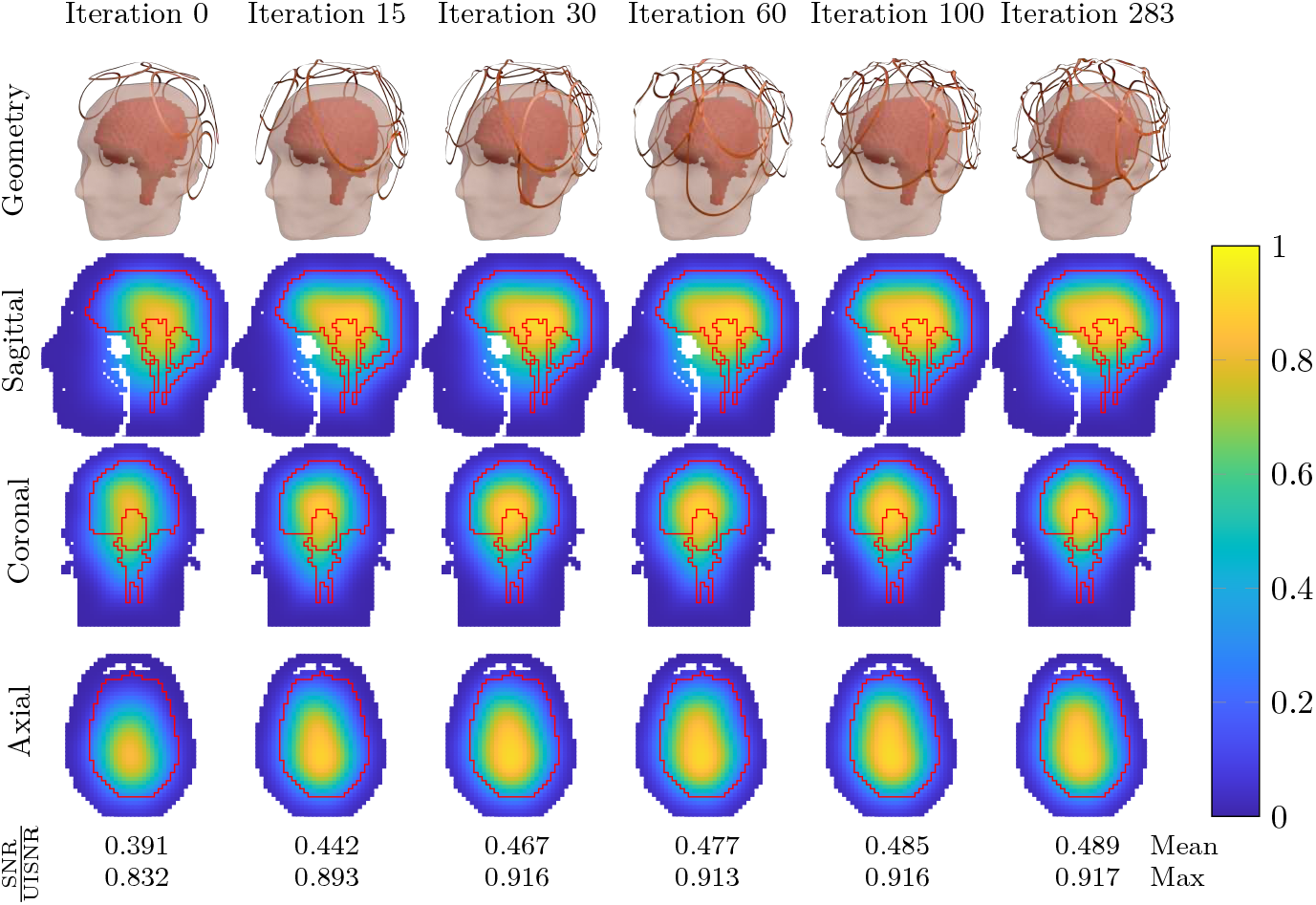
Evolution of the 12-coil geometry during the iterative optimization of the average SNR performance within the cerebrum ROI.

Figures 43 and 44 show the performance of the coil designs found at the completion of each optimization step (when *w*_1_ = 0.5, *w*_2_ = 0.5) in Figures 35 and 36 when Ella’s head model is used instead of Duke’s one. These results show that the optimized coil geometries at different optimization steps can also monotonically improve the SNR performance on Ella’s brain. The final optimized coil configuration increased the SNR performance in the center of Ella’s brain by 8% and improved the average SNR performance over the entire brain by 9%.

**Fig. 43:**
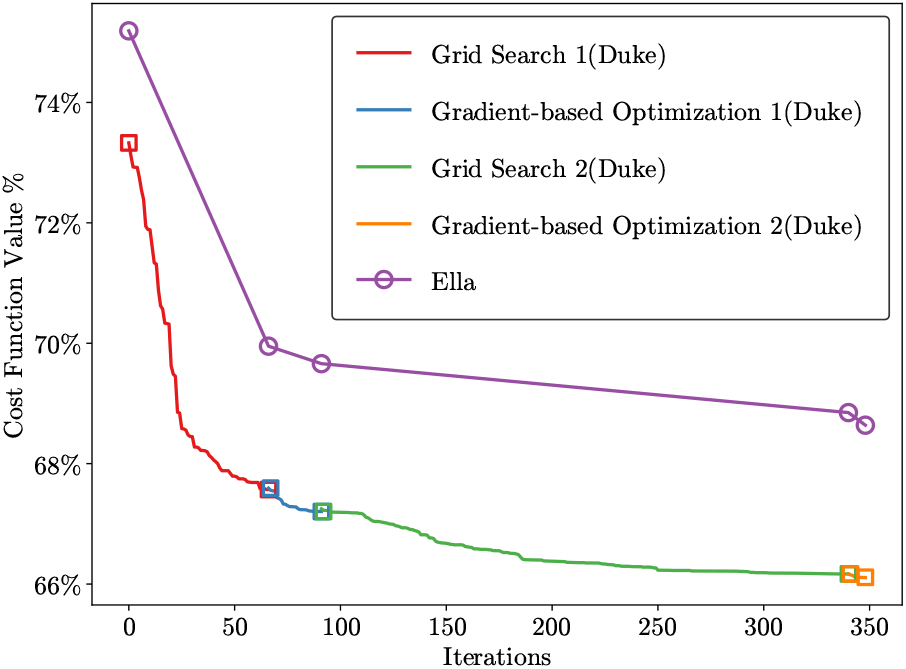
The cost function evolution for 12-coil optimization with the cerebellum and cerebrum ROIs using the Ella head model.

**Fig. 44:**
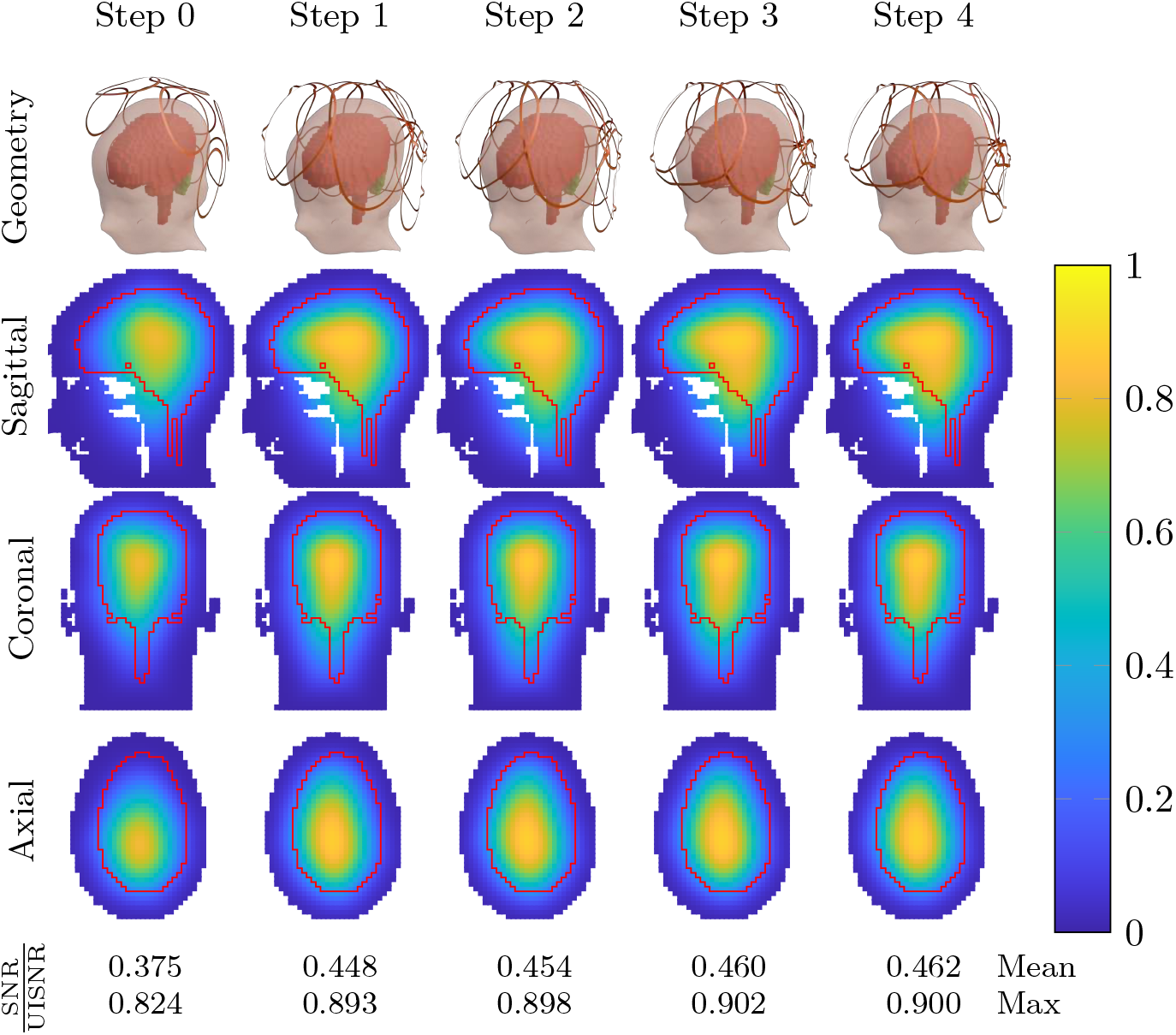
Evolution of the 12-coil geometry during the iterative optimization of the average SNR performance within the cerebellum and cerebrum ROIs using the Ella head model.

## 9 Discussion

The goal of this work was to develop an optimization framework for rational coil design for MRI applications. We developed an automatic tool that combines geometry processing techniques with state-of-the-art EM simulations to optimize coils using the UISNR as the reference. We eliminated user interactions, whereas setting up a single simulation could take days with the current coil design software. To achieve automation and computational efficiency, several innovations were introduced in this work. The new approach to solving the singular integrals during the assembly of the SIE operators considerably increases numerical precision, which is critical when these operators are used in iterative optimizations. We also introduced a novel C++ implementation that dramatically accelerates the assembly of the SIE operators. To perform iterative shape optimization of coil configurations, we needed to develop a rapid coil meshing method, including the automatic implementation of bridges to avoid contact between overlapping coil conductors, a feature not available in commercial coil design software. To efficiently simulate coils at each iteration, we developed an algorithm for automatic coil tuning and a method for ideal coil decoupling.

Previous work showed that ideal current patterns associated with the UISNR can provide a qualitative target to design coils that approach the optimum performance [7, 11]. Here, we have shown that certain coil configurations found by the optimization algorithm resemble the ideal current patterns, confirming graphically the near-optimality of common surface quadrature coil designs. Previous work on coil optimization cannot be directly compared with our results, since it was limited to one or two coils, used different design targets, and was performed in the quasi-static regime or using simplified analytical models. This is the first approach to use full-wave EM simulations and ultimate performance limits in the cost function to optimize coils for high-field MRI. As a proof of principle, in this work, we showed that it is computationally feasible to automatically optimize a 12-coil receive head array at 3 T using only grid search and gradient descent, achieving an absolute SNR performance in the brain region comparable to that of state-of-the-art 32-coil arrays [8]. The objective function that iteratively evaluates new coil geometries against the UISNR is smooth, which is favorable for gradient descent. However, optimizing denser arrays can become increasingly inefficient, so in future work we plan to calculate the gradients analytically using the Hermitian adjoint solution of the VSIE as in other optimization problems that utilize it [92, 93]. This will also allow us to broaden the design space parametrization, which in this work was limited to the size and position of the coils. We included various design constraints to ensure that the optimization would converge to coil configurations that could be practically constructed. Additional constraints could be added in the future to accommodate design requirements for particular clinical applications. Different constraints would also be needed to optimize transmit and transceive coils. For example, the ideal decoupling strategy developed for this work would not be feasible for transmitters since preamplifiers are not used during transmission. The objective function would also need to be modified to account for different metrics, such as the ultimate intrinsic SAR [6, 94] or the optimal transmit efficiency [4]. Note that the proposed optimization framework is flexible and generalizable enough to enable the incorporation of these changes in a straightforward manner.

The proposed optimization framework affects the entire flow of coil design and evaluation, and we expect that it will result in coils that yield higher performance than currently available coils. Since previous studies have shown that traditional coil designs cannot approach the ultimate SNR performance at ultra-high field MRI [9, 20, 39], this project could lead to the discovery of novel coil types that yield superior signal encoding capabilities. The coil designs generated by the automated pipeline would be directly importable into computer-aided design (CAD) environments for rapid prototyping, making application-specific, tailored coils a practical possibility.

This work aimed to introduce the new optimization framework. Several extensions will be pursued in future work, in addition to the already mentioned adjoint formulation and the implementation of other target optimization metrics. For example, replacing PWC with PWL basis functions would improve the accuracy of the UISNR [81]. However, the memory footprint and computational complexity would increase considerably, requiring the incorporation of novel strategies for tensor compression and efficient computation. We also plan to incorporate a wire integral equation solver to efficiently optimize coils with wire conductors, which are often preferred over flat copper strips to reduce coupling in arrays with many coil elements.

## 10 Conclusion

This work introduced the first fully automated software pipeline for optimizing coil design using ultimate performance limits as the reference benchmark. The pipeline integrates novel approaches for rapid EM simulations, shape optimization, and coil meshing. Several improvements and new features were incorporated into MARIE, an open-source VSIE solver tailored to MRI simulations. These include orders of magnitude higher precision in the assembly of the SIE operators, automatic coil tuning, and ideal decoupling of array elements. The coil design pipeline was demonstrated for optimizing head receive arrays with respect to the UISNR, for an increasing number of elements and different optimization target regions. The optimizations monotonically converged in all cases. The optimized 12-coil design at 3 T yielded approximately 10% higher SNR performance over an extended region of the brain compared to an arbitrary designed array.

## Supplementary information

Additional supplementary information may be found online in the Supplementary Information Section or at the end of this article.

## Acknowledgments

This work was supported in part by NIH R01 EB036483, NIH K99 EB035163, and NIH R01 EB024536, NSF 2313156, and was performed under the rubric of the Center for Advanced Imaging Innovation and Research (CAI^2^R, www.cai2r.net), an NIBIB National Center for Biomedical Imaging and Bioengineering (NIH P41 EB017183).

## Declarations

### Ethics approval and consent to participate

Not applicable

### Consent for publication

All authors have given their consent for publication.

### Data availability

Not applicable

### Materials availability

Not applicable

### Code availability

All code will be fully released as open-source on a dedicated repository after publication. Parts of the code used in this work for meshing and electromagnetic simulations are available at the following repositories:

https://github.com/thanospol/MARIE,

https://github.com/georgyguryev/MARIE_2.0,

https://github.com/qnzhou/nanospline

https://www.cs.cmu.edu/~quake/triangle.html

https://libigl.github.io/

### Author contribution

J.E.C.S. developed and implemented the new methods for automatic tuning, ideal decoupling, and improved SIE assembly. I.I.G. implemented the UISNR, ICP, and MRGF routines and integrated MARIE and MRGF to the optimization framework. S.W. developed and implemented the coil parametrization, coil meshing, and coil shape optimization routines. D.C. developed the graphical user interface. I.I.G., J.E.C.S., D.Zi., D.P., D.Zo., R.L., advised, analyzed data, and supervised the work. I.I.G., J.E.C.S., S.W., R.L., D.Zi., D.P., D.Zo., wrote the manuscript.

